# Function of Cytochrome P450 *CYP72A1182* in Metabolic Herbicide Resistance Evolution in *Amaranthus palmeri* Populations

**DOI:** 10.1101/2023.12.13.571468

**Authors:** Carlos Alberto Gonsiorkiewicz Rigon, Anita Küpper, Crystal Sparks, Jacob Montgomery, Falco Peter, Simon Schepp, Alejandro Perez-Jones, Patrick J. Tranel, Roland Beffa, Franck E. Dayan, Todd A. Gaines

## Abstract

Evolution of metabolic herbicide resistance is a major issue for weed management. Few genes and regulatory mechanisms have been identified, particularly in dicotyledonous weed species. We identified putative causal genes and regulatory mechanism for tembotrione-resistance in *Amaranthus palmeri*. Cytochrome P450 candidate genes were identified through RNA-seq analysis. We validated their functions using heterologous expression in *S. cerevisae*. Promoters of the candidate P450 genes were analyzed. We performed QTL mapping to identify genomic regions associated with resistance. *CYP72A1182* deactivated tembotrione. This gene had increased expression in other *A. palmeri* populations resistant to multiple herbicides, including tembotrione. Resistant plants exhibited polymorphisms in the promoter of *CYP72A1182*. We identified QTLs linked to herbicide resistance, including one on chromosome 4 approximately 3 Mb away from *CYP72A1182*. *CYP72A1182* is involved in tembotrione resistance in *A. palmeri*. Increased expression of this gene could be due to *cis*-regulation in the promoter, as well as *trans*-regulation from transcription factors. Further studies are in progress to test this hypothesis. The elucidation of regulatory genes is crucial for developing innovative weed management approaches and target-based novel molecules.

**HIGHLIGHTS:** Our study identifies that the *CYP72A1182* gene has a functional role in metabolic herbicide resistance in *Amaranthus palmeri* and is linked to *cis*-regulatory polymorphisms, advancing metabolic resistance understanding in dicots.

## INTRODUCTION

Evolution of herbicide resistance results from the selection pressure imposed by the frequent and consistent use of herbicides over time (Gaines et al. 2020). The emergence and spread of herbicide-resistant weeds poses a major challenge to sustainable agriculture, as they can diminish crop yields and increase the costs associated with weed control (Chauhan 2020; Oerke 2006). Herbicide resistance in plants can be selected through different mechanisms, primarily classified as target-site (TSR) and non-target-site resistance (NTSR) (Gaines et al. 2020; Rigon et al. 2020). Between these two groups, NTSR, specifically metabolic resistance, is a critical issue due to the potential interaction of a single metabolic resistance mechanism with herbicides spanning multiple modes of action (Rigon et al. 2020).

Plants possess a diverse array of enzymes involved in the detoxification of xenobiotics. Within these enzymes, cytochrome P450 enzymes (P450) play a key role in the initial step of the detoxification process (Rigon et al. 2020). P450s associated with herbicide metabolism have been identified from various crops (Brazier-Hicks et al. 2022; Kato et al. 2020; Pan et al. 2006; Pan et al. 2022; Siminszky et al. 1999) and implicated in evolved herbicide resistance in weeds. Notable examples from weeds include *CYP81A12*, *CYP81A21*, *CYP81A15*, and *CYP81A14* from *Echinochloa phyllopogon* (Dimaano et al. 2020); *CYP81A69* and *CYP81A70* from *Cynodon dactylon* (Zheng et al. 2022); *CYP81A10v7* from *Lolium rigidum* (Han et al. 2021) and *CYP77B34* from *Descurainia sophia* (Shen et al. 2022). Nearly all of the identified genes from weeds to date are from grass species (Rigon et al. 2020).

*Amaranthus palmeri* (Palmer amaranth) is a highly invasive and troublesome weed that poses a significant threat to agricultural crops and ecosystems. Native to the southwestern United States, this annual dicot weed has become a pervasive problem across North America and globally due to its aggressive growth, adaptability, and resistance to herbicides (Roberts and Florentine 2022; Ward et al. 2013). *A. palmeri* is dioecious, with separate male and female plants, obligate cross-pollination, and high within- and among-population genetic diversity (Gaines et al. 2021; Ward et al. 2013). Populations have evolved resistance to herbicide spanning nine different modes of action, including synthetic auxins (Foster and Steckel 2022; Shyam et al. 2022), glufosinate (Priess et al. 2022), and herbicides targeting acetolactate synthase (ALS) (Berger et al. 2016), 5-enolpyruvylshikimate-3-phosphate synthase (EPSPS) (Gaines et al. 2010), protoporphyrinogen oxidase (PPO) (Montgomery et al. 2021), photosystem II (PSII) (Jhala et al. 2014), 4-hydroxyphenylpyruvate dioxygenase (HPPD) (Küpper et al. 2018), very long chain fatty acid elongase (VLCFAE) (Hwang et al. 2023), tubulin biosynthesis (González-Torralva and Norsworthy 2021), and.

*A. palmeri* has evolved herbicide resistance through two main mechanisms, TSR and NTSR. The most commonly observed cases of resistance involve TSR to ALS- and EPSPS-inhibitors (Gaines et al. 2010; Palmieri et al. 2022). However, there are increasing resistance cases due to metabolic-based mechanisms (Küpper et al. 2018; Shyam et al. 2022), and further analysis is necessary to identify the specific genes involved and their regulation. RNA-seq analysis plays a crucial role in deciphering the genetic basis of herbicide resistance, especially when resistance is a multigenic trait (Gaines et al. 2014; Kohlhase et al. 2019). Additionally, with the availability of complete genomes for weed species (Montgomery et al. 2024), such as *A. palmeri*, conducting Quantitative Trait Locus (QTL) mapping experiments becomes a valuable approach for determining the genetic architecture of resistance (Murphy et al. 2021). By combining comprehensive gene expression data obtained from RNA-seq analysis with QTL mapping, researchers can gain valuable insights into the genetic factors underlying resistance and other important characteristics in weeds.

This research aimed to identify and functionally validate metabolic genes associated with resistance to tembotrione (an HPPD inhibitor) within a particular population of *A. palmeri*. We subsequently sought to determine whether an identified gene, *CYP72A1182*, was also associated with resistance in other suspected tembotrione-resistant *A. palmeri* populations. The research also sequenced the promoter and ran a QTL analysis to investigate potential regulation of *CYP72A1182*.

## MATERIALS AND METHODS

### Plant material for QTL mapping and RNA-seq analysis

The resistant (NER) and susceptible (NES) *A. palmeri* populations were collected from fields in Shickley, Nebraska in 2011. NER is resistant to atrazine and the HPPD inhibitors tembotrione, mesotrione, and topramezone (Jhala et al. 2014; Küpper et al. 2018). The enhanced metabolism of tembotrione to hydroxy-tembotrione was identified as the mechanism of herbicide resistance in the NER population (Küpper et al. 2018). To minimize the impact of genetic differences unrelated to metabolic herbicide resistance, QTL mapping and RNA-seq was performed on pseudo-F2 plants generated from controlled pairings of NER and NES parents. The approaches for the generation of the pseudo-F2 plants are in Methods **S1** and Fig. **S1**.

### RNA-sequencing analysis

Descriptions of plant population, herbicide application, plant tissue collection, and library preparation are in Methods **S1**. Raw reads were pre-processed by removal of library adapter sequences, removal of low quality reads, and assessing the quality control using fastp (Chen et al. 2018). Read alignment of the 108 libraries to the available male genome of *A. palmeri* (v1.1, id55760) (Montgomery et al. 2020) was performed using Hisat2.2.1 (Kim et al. 2019). Most of the reads (>77%) aligned concordantly once, while >7% aligned concordantly more than one time and about 14% of reads aligned not concordantly. Mapped reads were assigned to gene features using featureCounts (Liao et al. 2014).

### P450 gene sequences and phylogenetic tree

Consensus sequences of the candidate P450 genes were extracted from the transcriptome and aligned to the reference *A. palmeri* genome (Montgomery et al. 2020) to analyze single nucleotide polymorphisms (SNPs). PCR was performed to amplify the coding sequences in four each of sensitive (S) and resistant (R) plants from the pseudo-F2 population that were used for the RNA-seq experiment. cDNA synthesis was performed using ProtoScript® II First Strand cDNA synthesis kit with 1 µg of RNA. Protein sequences from cytochrome P450 known to metabolize herbicides in different plant species were obtained from NCBI. Multiple protein alignment was performed using ClustalO and used for tree construction with Neighbor-joining method using Geneious Prime® 2023.0.1.

### Quantitative reverse transcription-PCR (RT-qPCR) validation

NES and NER populations were grown and tembotrione herbicide was applied at 91 g a.i. ha^−1^ as previously described. Four plants of each population were used for gene expression analysis validation. The youngest leaf tissue was collected at 0, 3, 6, and 12 h after herbicide application (HAT) in 2 mL Eppendorf tubes and placed in liquid nitrogen. The tubes were kept at -80 °C for further analysis. Tissue was ground with 3 mm stainless steel beads in a TissueLyser (Qiagen) with intensity of 30 for 1 min. RNA was isolated using Direct-zol RNA Miniprep from Zymo Research. cDNA synthesis was performed using ProtoScript® II First Strand cDNA synthesis kit using 1 µg of RNA and purified with DNase I.

Relative gene expression was analyzed on a T100 Thermal Cycler (BioRad), using SsoAdvanced™ universal SYBR® Green supermix. Reaction mixtures consisted of 10 µL of SsoAdvanced universal SYBR® Green supermix (2X), 2.5 µL of forward and reverse primers at 10 µM, and 5 µL of cDNA (1:20 dilution). Thermocycler conditions consisted of an initial step of 30 sec at 95°C followed by 35 cycles of 5s at 94°C and 30 s at 60°C. Melt curve analysis was added using 65°C with 0.5°C increments of 5 s/step.

The reference genes used were *18S rRNA* and *Actin7*. These two genes had the best gene stability as assessed by the NormFinder algorithm (Andersen et al. 2004) (Tab. **S1**). Primers for the reference genes were designed based on conserved regions after alignment of *18S (FJ669720.1), actin* (HQ656028.1), and *TUB* (XM_010693569.3) from *Beta vulgaris* with the reference genome from *A. palmeri* (CoGe – v1.1, id55760). The candidate genes tested based on RNA-seq results were *CYP72A1182*, *CYP72A1027*, *CYP72A1015*, and *CYP81CJ2*. Another *CYP72A-like* gene, hereafter named as *CYP72A*_*4286* was used as a cytochrome P450 not associated with herbicide resistance based on RNA-seq data. Primers for candidate genes were designed on regions based on consensus sequences from the transcriptome and Sanger sequencing of the genes and are listed in Tab. **S2**.

### Heterologous expression of CYPs

The genes of interest were transformed and expressed in yeast for further investigation of their role in tembotrione metabolism. Coding sequences of the candidate genes were synthesized, optimized for yeast transformation, and inserted in the pUC-WG/ampv vector by GENEWIZ (Azenta Life Sciences). The gene sequences used were *CYP72A1182* allele 1 (GenBank accession number: OR596705) and 2 (OR596706), *CYP72A1027* (OR596707), *CYP72A1015* allele 1 (OR596708) and 2 (OR596709), and *CYP81CJ2* allele 1 (OR596710) and 2 (OR596711). As a positive control, the gene *CYP81A9* from corn, known as *Nsf1*, was used. This gene was identified as the major resistance locus in corn for the herbicides nicosulfuron and mesotrione (Williams and Pataky 2008). Restriction sites for BamHI and EcoRI were added to the 5’ and 3’ ends, respectively. Kozak sequence (AAAAAATCT) was added at the 5’ end as a protein translation initiation site. Restriction enzyme reactions were performed to cut the optimized sequence from the vector using 1 μL EcoRI, 1 μL BamHI, 500 ng of the vector, with incubation at 37°C for 2 h. Reaction products were run on 1% agarose gel electrophoresis and the gene band was isolated and purified using a DNA Gel Purification Kit from New England Biolabs®.

The pYES2 yeast expression vector was used for recombinant expression. It contains the *URA3* gene for selection in yeast and 2µ origin for high-copy maintenance and GAL1 promoter to express protein (Thermo Fisher Scientific). The ligation reaction was performed using LigaFast™ Rapid DNA Ligation System from Promega. The reaction consisted of 50 ng digested pYES2 plasmid vector, 5 μL 2X Rapid Ligation Buffer, 1 μL T4 DNA ligase (3 Weiss unit/μL), 50 ng of gene, and purified water up to 10 µL. The reaction was incubated overnight at 4°C. The product from the reaction was used to transform *E. coli* using the One Shot® TOP 10 kit (Invitrogen). The transformed cells were plated with ampicillin (100 μg/mL) and incubated at 37°C overnight. Single colonies were selected, and gene insertion was confirmed by colony PCR using high-fidelity PrimerStar HS DNA polymerase. The gene sequence was confirmed by Sanger sequencing. Plasmids were isolated using ZymoPure II Plasmid Miniprep Kit.

### Yeast transformation and tembotrione incubation

Yeast transformation protocol is described in Methods **S2**. Single colonies containing empty pYES2 or the candidate CYP genes were incubated in 15 mL 2% raffinose medium and allowed to grow for 2-3 d at 30 °C. Cell density was measured spectrophotometrically until the OD600 in 20 mL of induction medium was 5. The volume was removed and pelleted at 1,500 X g for 15 min at 4 °C. The cells were resuspended with 1 mL of induction medium containing 2% galactose and inoculated into 20 mL of the same medium. Tembotrione at 1500 μM diluted in EtOH was applied right after the cells were incubated in the induction medium. The yeast cells were incubated at 30°C at 200 rpm. Twenty-four hours after herbicide application, 5 mL of the medium were collected in 15 mL falcon tubes and cells were pelleted at 1,500 X g for 5 min. The supernatant was collected, and cleaned by adding 5 mL of 5% acetic acid acetonitrile + QuEChERS, vortexed for 30 s, and centrifuged for 15 min at 2,000 X g. The supernatant was collected, filtered through a nylon 13 mm by 0.2 µm (Econofltr Nyln, Agilent Technologies), and injected in the LC-MS/MS.

### LC-MS/MS protocol

LC-MS/MS system consisted of a Nexera X2 UPLC with 2 LC-30AD pumps, a SIL-30AC MP autosampler, a DGU-20A5 Prominence degasser, a CTO-30A column oven, and SPD-M30A diode array detector coupled to an 8040 quadrupole mass-spectrometer. For tembotrione, the MS was in negative mode with an MRM optimized for 439.1>226.05 and set for 100 ms dwell time with a Q1 pre-bias of 11.0V, a collision energy of 11.0V and a Q3 pre-bias of 14.0V. For hydroxy-tembotrione, the MS was in negative mode with an MRM optimized for 455.1>419.05 and set for 100 ms dwell time with a Q1 pre-bias of 11.0V, a collision energy of 11.0V and a Q3 pre-bias of 14.0V. The samples were chromatographed on a 100X4.6 mm Phenomenex Kinetex 2.6 μm biphenyl column maintained at 40°C. Solvent A consisted of water with 0.1% formic acid and solvent B was acetonitrile with 0.1% formic acid. The solvent program started at 80% B and increased to 100% B in 3.5 min and maintained at 100% for 2 min. The solvent was returned to 80% B and maintained there for 3 min before the next injection. The flow rate was set at 0.4 mL/min and samples were analyzed as 1 μL injection volumes.

### *CYP72A1182* and *CYP81CJ2* promoter amplification

Pseudo-F2 plants were grown in the greenhouse and the youngest leaf tissue was collected when the plants reached four to five true leaf stage. Tembotrione was applied at 77 g ha^−1^ rate.

Five each of R and S plants from a pseudo-F2 population were chosen for promoter amplification. DNA extraction was performed using hexadecyltrimethylammonium bromide (CTAB) method (Doyle and Doyle 1990). DNA quantification was carried out using Nanodrop 2000c (Thermo Fisher Scienfitic). Primers were designed by Primer3Plus (https://www.bioinformatics.nl/) using as reference the available draft genome of *A. palmeri* (Montgomery et al. 2020). The primer sequences used are listed in Tab. **S2**. The methods used to amplify the gene promoter are available in Methos **S3**.

### Involvement of *CYP72A1182* in different tembotrione-resistant *A. palmeri* populations

Seeds from *A. palmeri* populations were collected from agricultural fields in 2019 in the United States with suspected herbicide resistance to mesotrione and tembotrione. Whole-plant dose response, ^14^C-tembotrione metabolism, P450 gene expression and gene copy number were evaluated to test the hypothesis that the same candidate P450 gene is involved in the resistance mechanism in these additional populations. The methods are described in Methods **S4**.

### Mapping of tembotrione resistance in NER *A. palmeri* population

#### Library preparation and QTL identification

Dose-response and segregation analysis methods are described in Methods **S5**. Plant tissue was collected before herbicide application for DNA extraction from single leaves following a modified CTAB method. DNA samples were assessed for quality using Qubit. Double-digest restriction site-associated DNA sequencing (ddRADseq) libraries were generated with ApeKI and sequenced using NovaSeq S4 with 150 bp paired-end reads (Illumina). The average yield was around 7.4 M reads per library and the mean quality scores were over Q30 for all libraries. The sequencing was performed at the University of Minnesota Genomics Center.

The ddRAD-seq libraries were trimmed using trimmomatic v0.36 (Bolger et al. 2014), aligned to the reference male genome of *A. palmeri* (scaffold file - v1.1, id55760) (Montgomery et al. 2020) with Burrows-Wheeler Aligner (Li and Durbin 2009) and variants called using GATK 4.2.0 (Poplin et al. 2018). Samples were computationally binned by shoot fresh mass. The variant sites were separated by SNPs and insertion/indels and hard filter was performed as following for SNPs - QD < 2.0, QUAL < 30.0, SOR > 4.0, FS > 20.0, MQ < 50.0 and insertion/indels QD < 2.0, QUAL < 30.0, FS > 200.0. The variant sites were filtered based on depth (at least five reads) and only variants that were homozygous were selected. R/qtl2 package was used to find QTLs (Broman et al. 2019). The analysis was performed separately for two distinct crosses (A and B) and combined A + B. QTL intervals were calculated using Bayes credible intervals with the function bayes_int(). To determine the threshold for identifying potential QTL, 1000 permutation tests were conducted at a confidence level of 95%. The critical F value obtained from this analysis was used as the criterion to declare putative QTL. The functional analysis of the genes present in QTLs was conducted using DAVID Bioinformatics Resources v6.8 (Sherman et al. 2022). Statistical analysis for all the experiments is described in Methods S6.

## RESULTS

### Differentially expressed genes (DEGs)

A total of 20,846 genes were analyzed for differential expression out of the initial set of 29,758 genes assigned by featureCounts. Gene-wise dispersion curve was fitted using the DESeq2 model (Fig. **S2**). There was favorable dispersion pattern, with decreasing dispersion as the mean expression levels increased, indicating a good fit of the DESeq2 model to the analyzed data. The principal component analysis (PCA) plot revealed that PCA1 and PCA2 accounted for 35 and 21% of the variation in the data, respectively (Fig. **S3**). PCA1 separated cross A from cross B, while PCA2 exhibited separation of gene clusters based on HAT within each cross. When clustering all 72 analyzed transcriptome samples, two major clusters emerged, with samples from different crosses grouped together, indicating similarities in gene expression response (Fig. **S4**).

Differentially expressed genes (DEGs) were observed in different comparisons. In the contrast comparison of R versus S at constitutive (0 HAT), 6 HAT, and 12 HAT after herbicide application, R had 37, 2, and 9 up-regulated genes and 3, 2, and 4 down-regulated genes, respectively (Fig. **1a**). Among these genes, only two genes were commonly up-regulated (Fig. **1a**). For the analysis of the genes that were responsive to herbicide application for each phenotype, 39, 530, and 106 genes were up-regulated in S and 6, 185, and 160 were up-regulated in R for the comparisons of time 6 vs 0 HAT, 12 vs 0 HAT, and 12 vs 6 HAT, respectively (Fig. **1a**). The heatmap illustrates the expression of DEGs from these comparisons and indicates a separation in clusters of resistant and susceptible plants (Fig. **1a**).

**Fig. 1.**
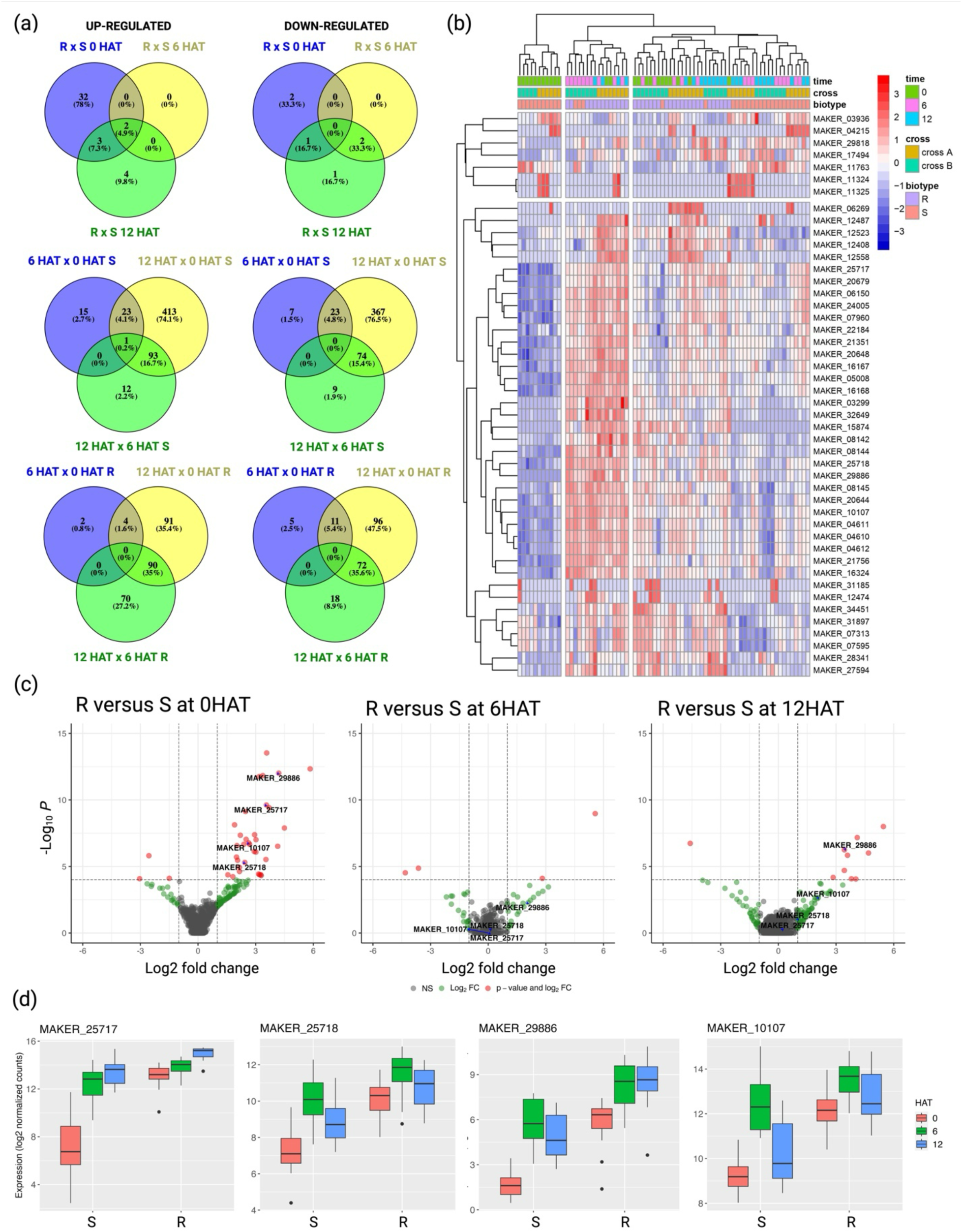
Analysis of differentially expressed genes (DEGs) in Pseudo-F2 *Amaranthus palmeri* susceptible (S) and HPPD-resistant (R). (a) The Venn diagram of the up- and down-regulated DEGs indicating unique and common DEGs for 18 different comparisons. (b) Heatmap of top DEGs for the comparisons R vs S at 0, 6 and 12 h after herbicide treatment (HAT). (c) Volcano plots displaying gene expression differences between *Amaranthus palmeri* and treatments of contrast of R versus S before herbicide application (0 HAT), contrast of R versus S at 6 HAT, and contrast of R versus S at 12 HAT. Differentially expressed genes with significant thresholds of p-value <0.0001, adjusted p-value <0.05, log2 fold-change >1 or < -1 are represented by red circles. Dashed lines represent significance thresholds of adjusted p-value<0.05 and log2 fold-change >1 or < -1. (d) The log2 normalized counts of top four cytochrome P450 genes in response to tembotrione in S and R population, MAKER_25717 (*CYP72A1182*), MAKER_25718 (*CYP72A1027*), MAKER_29886 (*CYP72A1015*), and MAKER_10107 (*CYP81CJ2*). Differently expressed genes had a threshold of adjusted p-value of 0.05 and log2 fold-change 1.

Several genes associated with the detoxification of xenobiotic substances were consistently identified, suggesting an enhanced genetic metabolism in plants with resistance. These genes included four cytochrome P450 genes, four glutathione-S-transferase genes, seven glycosyltransferase genes, a disease resistance protein, and a detoxification 27 gene (Tab. **S3**). Additionally, two MADS-box transcription factors of types 23 and 27 were found to be up-regulated and arise hypothesis of gene regulation in resistant plants. The resistance mechanism in the resistant population was characterized by increased herbicide metabolism including an initial hydroxylation of the tembotrione molecule (Küpper et al. 2018); therefore, among all differentially expressed genes (DEGs), our focus was on the four cytochrome P450 genes that were consistently up-regulated in the R plants. These genes had the genomic IDs MAKER_29886, MAKER_25717, MAKER_10107, and MAKER_25718, and exhibited fold changes of 18.3, 11.8, 6.2, and 5.4, respectively (Fig**. 1b-d** and Tab. **S3**). None of these genes had differential expression at the 6 HAT time point compared to the susceptible plants (Fig. **1c**), and only one gene (MAKER_29886) exhibited a reactivation at the 12 HAT (Fig. **1c**). This indicates that the susceptible plants responded to the herbicide treatment (Fig. **1c**), where an increased transcription of these genes was observed at 6 and 12 HAT. However, the resistant plants constitutively expressed these cytochrome P450 genes at a higher level prior to herbicide treatment.

Constitutively expressed P450s, MAKER_25717, MAKER_25718, and MAKER_29886, had significant similarity to *CYP72A219* from *Spinacia oleracea* (NCBI: XM_056843192.1), with identities of 76.2, 74.3, and 76.3% respectively. Similarly, the gene ID MAKER_10107 assigned the name *CYP81CJ2* had high similarity (73.3%) to *CYP81E8* from *Chenopodium quinoa* (NCBI: XP_021724107.1). Hereafter, the genes MAKER_25717, MAKER_25718, MAKER_29886, and MAKER_10107 will be referred to as *CYP72A1182*, *CYP72A1027*, *CYP72A1015*, and *CYP81CJ2* respectively, based on P450 gene annotation and naming from a recently assembled *A. palmeri* genome (Raiyemo et al. 2024).

### SNPs in cytochrome P450 candidate

There were no single nucleotide polymorphisms (SNPs) causing amino acid changes that were consistently different between S or R individuals in the P450 genes. *CYP72A1182* has a coding sequence of 1544 bp and 517 amino acids. *CYP72A1027* has a coding sequence of 1578 bp and 526 amino acids. Both R and S phenotypes shared the same two alleles for these genes. *CYP72A1015* has a length of 1394 bp and 464 amino acids. Susceptible plants had a single allele while R plants had the same allele and an additional allele with amino acid substitutions of Thr307Ser, Phe388Leu, and Ile407Val. *CYP72A1182* and *CYP72A1027* are localized on chromosome four and *CYP72A1015* is localized in chromosome 16. *CYP72A1182* shares identity with *CYP72A1015* and *CYP72A1015* of 76.1 and 55.1%, respectively. Two alleles of *CYP81CJ2* were present in both S and R samples. The gene has a length of 1470 bp and 1479 bp in each allele. The short allele has a deletion of 18 bp in position 35, and the longest allele has a deletion of 9 bp in position 37 when aligned to the draft genome. In the available *A. palmeri* genome, this specific region of the gene exhibits a triplication of PPS amino acids, which may be due to incorrect assembly. All cytochrome P450 possess a conserved cluster of proline in the membrane hinge. Similarly, the sequenced cytochrome P450 exhibits this conserved region.

In the phylogenetic tree analysis, *CYP72A* genes from *A. palmeri* clustered with *CYP72A31* from *Oryza sativa*, which can metabolize bispyribac-sodium and bensulfuron-methyl (Saika et al. 2014). *CYP81CJ2* clustered with other P450 of family 81 from grasses but with some significant distances between them (Fig. **S5**). This gene has a low identity when aligned with the other P450s. *ApCYP81CJ2* shares identity of 40.5%, 41.0%, and 39% with *ZmCYP81A9* (Genbank: EU955910.1), *LrCYP81A10v7* (Genbank: MK629521.1), and *EpCYP81A12* (Genbank:AB818460.1), respectively, known to metabolize herbicides in grasses (Brazier et al. 2002; Brazier-Hicks et al. 2022; Han et al. 2021; Iwakami et al. 2014).

### P450 qPCR validation

*CYP72A1182*, *CYP72A1015*, and *CYP72A1015* were upregulated in NER plants at 3, 6, and 12 HAT, respectively (Fig. **S6**). Among the *CYP72A* genes, *CYP72A1182* had a higher up-regulation at 3 and 6 HAT. Additionally, *CYP81CJ2*, located on chromosome 4, had up-regulation specifically at 6 HAT (Fig. **S6**). These observations deviate from those obtained in the RNA-seq analysis, which indicated that these genes are constitutively higher expressed in resistant plants (Fig. **1**). However, this discrepancy arises due to the utilization of the parental populations NES and NER for gene validation, which exhibit significant individual-level variation in their response to tembotrione. Nevertheless, despite the disparity, the consistency of these findings indicates that the up-regulation of these genes in NER plants differs from that in NES plants.

### *CYP72A1182* metabolizes tembotrione in yeast

Following a 24 h incubation period with tembotrione, some transformed yeast treatments converted tembotrione into hydroxy-tembotrione. The chromatogram of the empty vector pYES2 revealed solely the presence of the parent tembotrione peak at a retention time of 2.9 min (Fig. **2**). Notably, the *Nsf1* gene exhibited a pronounced affinity for tembotrione, producing hydroxy-tembotrione peak at a retention time of 2.7 min and a dihydroxy tembotrione peak at 2.5 min (Fig. **2**). Among all the candidate P450 genes tested, only *CYP72A1182* converted the parent tembotrione into hydroxy-tembotrione. Both alleles of this gene in the WAT11 and WAT21 strains metabolized the herbicide. Conversely, *CYP72A1015*, *CYP72A1015*, and *CYP81CJ2* did not exhibit any ability to metabolize tembotrione. Based on these results, it can be concluded that of the candidate P450 genes, only *CYP72A1182* possesses the capacity to recognize the tembotrione molecule and effectively metabolize it.

**Fig. 2.**
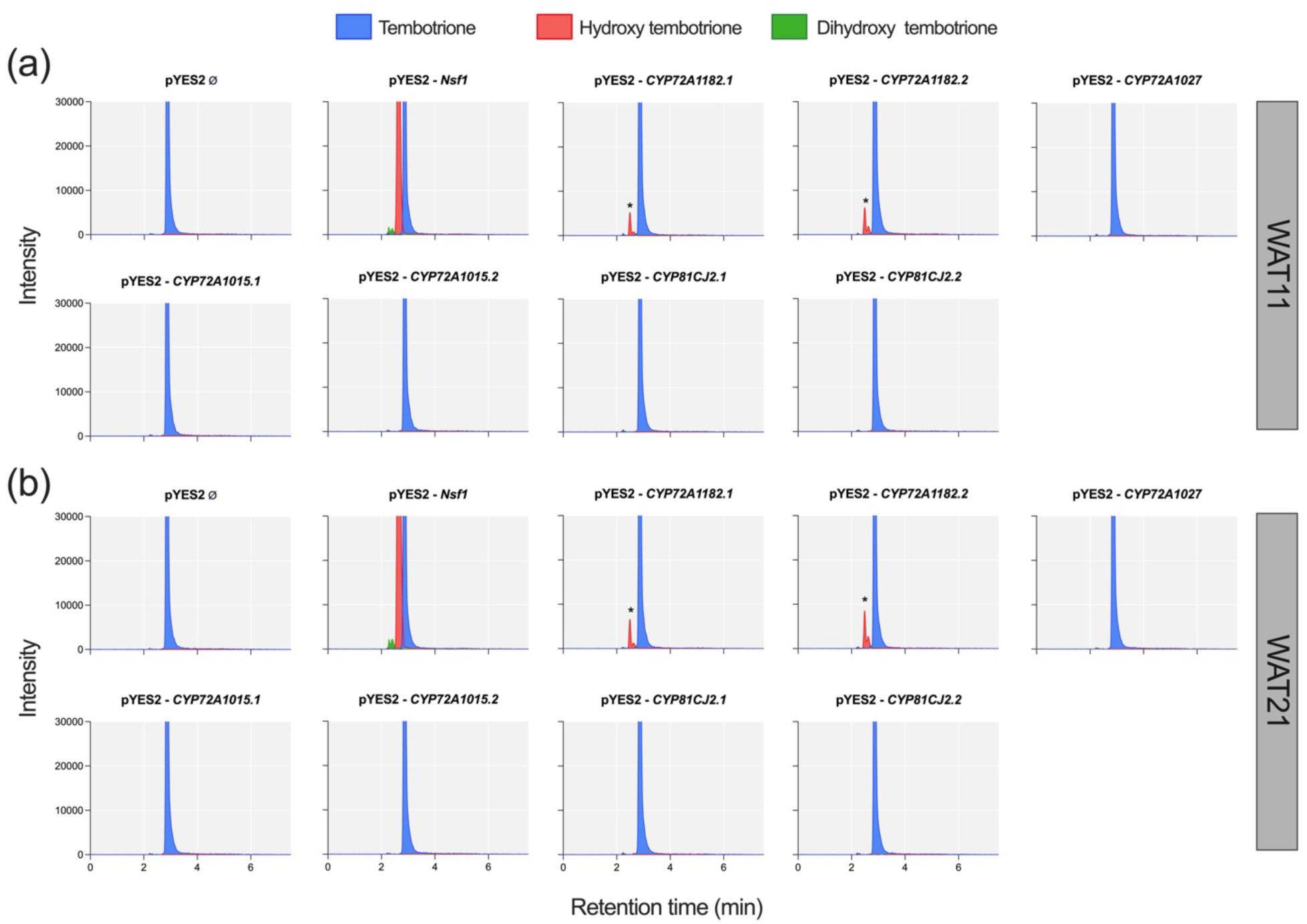
LC-MS/MS chromatogram of metabolites formed in yeast transformed with P450 genes from *A. palmeri*. Blue peak – parental tembotrione, read peak – hydroxy-tembotrione, green peak, dihydroxy-tembotrione. (a) WAT11 and (b) WAT21 yeast strain carrying *Arabidopsis thaliana* cytochrome P450 reductase 1 and 2, respectively. * at least 10-fold higher peak than the background signal.

### CYPs promoter analysis

The DNA segment amplified upstream of *CYP81CJ2* was 1,250 base pairs for both the sensitive and resistant plants (Fig. **3**). Pairwise sequence alignment between two samples revealed 100% identity between the promoters of S and R (Fig. **3a** and Notes **S1**). For the promoter of the *CYP72A1182* gene on chromosome four (MAKER-25717), the primers amplified regions ranging from 1991 to 2004 base pairs for S plants and from 1747 to 1768 base pairs for R plants. Pairwise alignment of *CYP72A1182* promoters revealed sequence differences between S and R, with identity of 76.8% (Notes **S1**). Resistant plants had unique insertions, such as a cytosine at position 45 bp, an 8 bp insertion at position 458 bp, an 8 bp insertion at position 1247 bp, and a 31 bp insertion at position 1291 bp upstream. Sensitive plants, on the other hand, had unique insertions, including a 13 bp insertion at position 82, a 10 bp insertion at position 309 bp, and a 277 bp insertion at position 746 bp upstream of the gene (Fig. **3b** and Notes **S1**).

**Fig. 3.**
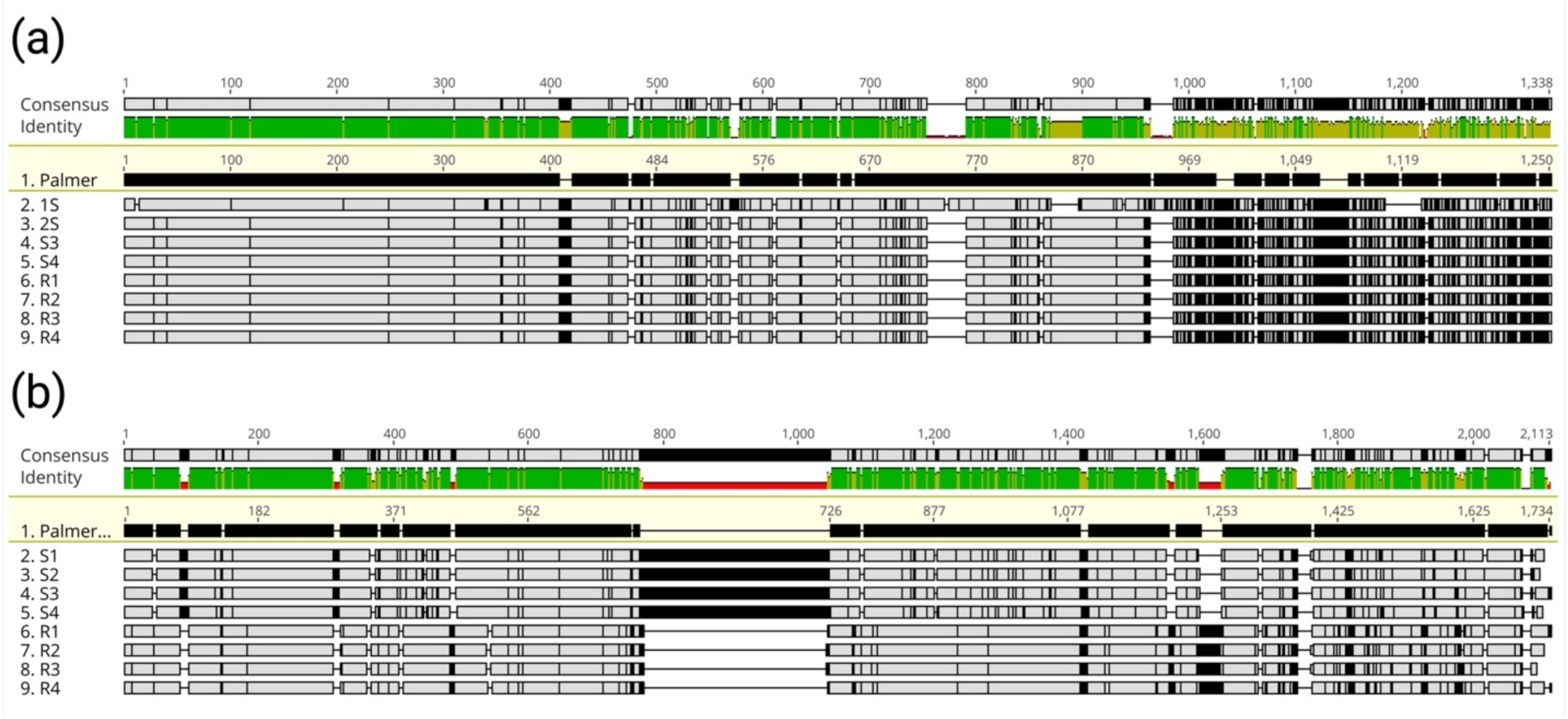
MUSCLE v5 multiple alignment of *CYP81CJ2* (a) and *CYP72A1182* (b) promoter sequences. The promoter sequences were reverse-oriented from the reference genome sequence to enable viewing in + strand orientation. Black space indicates differences from the reference *A. palmeri* genome (id 55760).

A total of fifteen significant binding-site motifs were identified in the promoter region of *CYP81CJ2*. Interestingly, all motifs were found in both S and R promoters (Fig. **S7**). In the case of the *CYP72A1182* promoter, a total of twenty-five motifs were discovered. Among these, two motifs were specific to the S promoter, two were specific to the R promoter, and twenty-one motifs were shared between S and R (Fig. **S7** and **S8**). Notably, the R promoter exhibited replication of four motifs. Some motifs were identified as a binding site for transcription factors such as MYB88 (AT2G02820), suppressor factor AIF1 (AT3G05800), and abiotic stress-responsive transcription factor DREB (AT1G77200) based on GOMo analysis (Fig. **S8**). Furthermore, the motif GTTATTTAGTAACTTWKYGTD, which serves as a binding site for the MYB gene transcription factor, was triplicated in the R promoter (Fig. **S8**).

### Whole-plant dose response, metabolism, and *CYP* expression in different herbicide resistant *A. palmeri*

Based on the whole-plant dose response (Fig. **S9**), the confirmation of *A. palmeri* populations with suspected tembotrione resistance revealed distinct variations, with resistance index ranging from 2.4 to 7.9 (Tab. **S4** and **S5**). Tembotrione metabolism was assessed in the *A. palmeri* populations using ^14^C herbicide (Fig. **S10**). The method used in the present study was different from the previous study (Küpper et al. 2018), hence, retention times are slightly different. Parental tembotrione had a retention time of 66.9 min and its major metabolites had retention times of 47.3 (M1), 50.1 (M2), 56.3 (M3), 59.0 (M4), and 67.7 min (M5) (Fig. S11 and S12). NER had higher % area peak of hydroxy-tembotrione (M3/M4), the main metabolite associated with herbicide resistance (Küpper et al. 2018), than the sensitive population at 12 HAT (Fig **S10**). The other resistant populations were similar to NER, except for WR2019-044 and WR2019-273 (Fig. **S10**). The faster hydroxy-tembotrione production in most populations suggests that enhanced metabolism capacity is the main mechanism of HPPD-resistance of these populations. *CYP72A1182* was upregulated in most resistant populations, either before treatment or 6 HAT (Fig. **S1**). The populations WR2019-034, -137, -140, -199, and -274 had a significantly higher relative *CYP72A1182* expression compared to the sensitive population, ranging from 100 to 658-fold increase. In contrast, the expression of *CYP81CJ2* was not constitutively higher or responsive to tembotrione treatment in *A. palmeri* populations, except in WR2019-140 (Fig. **S1**). The gene copy number of *CYP72A1182* was similar across all resistant populations and with that of sensitive plants (Fig. **S14**). The higher gene expression of *CYP72A1182*, constitutively and after tembotrione application in resistant plants, suggests a crucial role in herbicide resistance in the newly characterized tembotrione-resistant populations.

### Degree of dominance and QTL identification

The ED_50_ of NES and NER were 7.9 and 37.3 g, respectively (Tab. **1**), lower than previously reported for these populations (Küpper et al. 2018). However, the resistance index (RI) between these two populations was similar, 4 compared with 3.3 from the previous study (Küpper et al. 2018). Both F1 populations were intermediate between NES and NER in response to tembotrione (Fig. **4**). The degree of dominance was calculated and indicated a co-dominant or semi-dominant trait (cross A = 0.42 and cross B = 0.67) based on the formula proposed by Bourguet and Raymond (1998). Pseudo-F2 populations, resulting from the cross between A and B, exhibited survival rates of 11.4% and 8.8%, respectively. These survival rates are similar to the parental homozygous line NER (12.4%), which was cultivated alongside the pseudo-F2 plants.

**Tab. 1.**
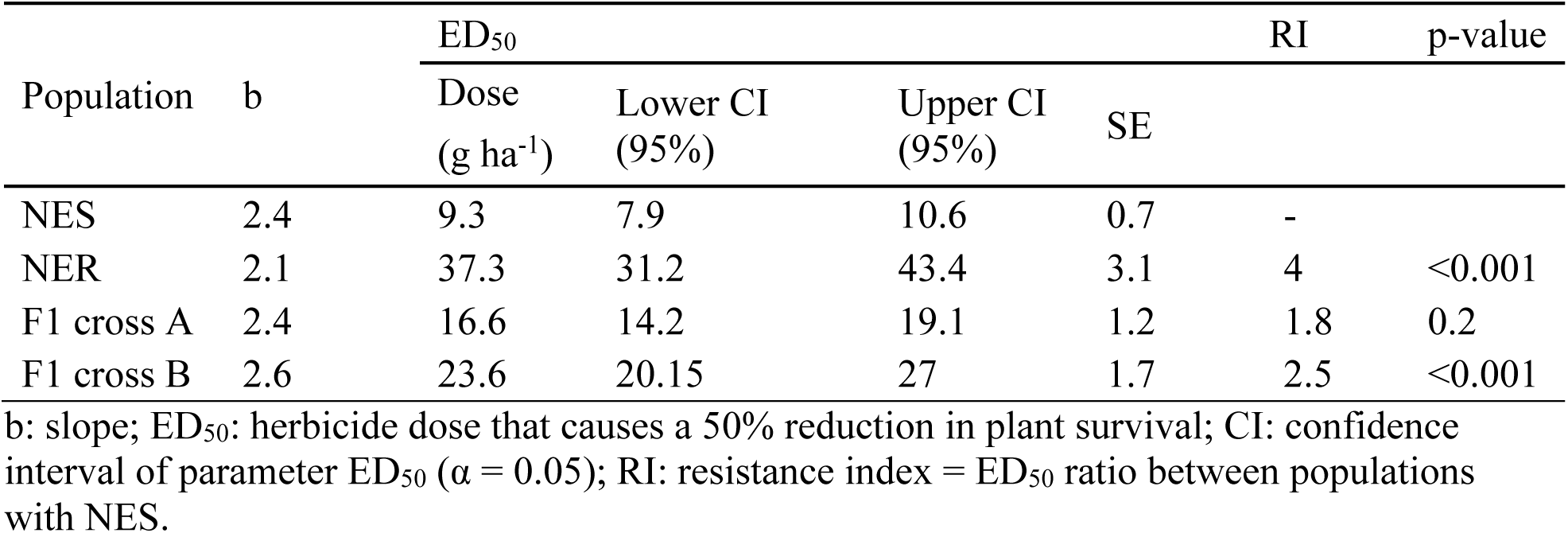
Logistic equation parameters and resistance index (RI) for survival (% of untreated control) in parental *Amaranthus palmeri* herbicide-susceptible (NES), herbicide-resistant (NER), F1 cross A, and F1 cross B populations in response to tembotrione.

| Population | b | ED<sub>50</sub> |  |  | SE | RI | p-value |
| --- | --- | --- | --- | --- | --- | --- | --- |
|  |  | Dose<br>(g ha<sup>-1</sup>) | Lower CI<br>(95%) | Upper CI<br>(95%) |  |  |  |
| NES | 2.4 | 9.3 | 7.9 | 10.6 | 0.7 | - |  |
| NER | 2.1 | 37.3 | 31.2 | 43.4 | 3.1 | 4 | <0.001 |
| F1 cross A | 2.4 | 16.6 | 14.2 | 19.1 | 1.2 | 1.8 | 0.2 |
| F1 cross B | 2.6 | 23.6 | 20.15 | 27 | 1.7 | 2.5 | <0.001 |

b: slope; ED<sub>50</sub>: herbicide dose that causes a 50% reduction in plant survival; CI: confidence interval of parameter ED<sub>50</sub> ( $\alpha = 0.05$ ); RI: resistance index = ED<sub>50</sub> ratio between populations with NES.

**Fig. 4.**
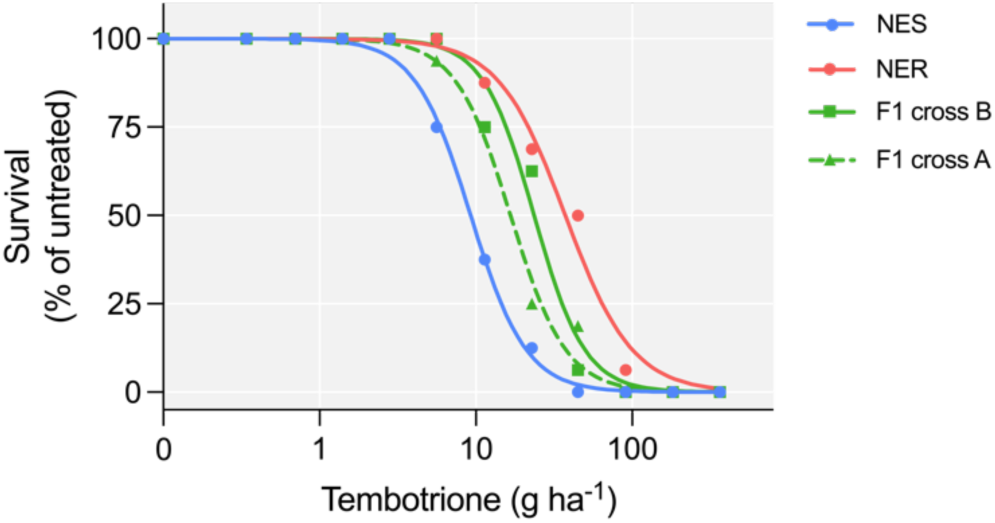
Dose-response of *Amaranthus palmeri* herbicide susceptible (NES), resistant (NER), F1 cross A and F1 cross B plants in response to tembotrione.

Frequency distribution of fresh weight after herbicide treatment on pseudo-F2 plants had two distinct peaks for each cross, with one peak each indicating phenotypic similarity with each parental population. A normal distribution of R plants had higher biomass (red bars) and the same was identified for S samples (Fig. **5a**). The variant calling analysis identified over 3.4 million SNP sites in the *A. palmeri* genome. After filtering, a total of 4,404 SNP variants were detected throughout the entire *A. palmeri* genome. Genome scanning in cross A did not detect any QTLs surpassing the LOD threshold of 6.4 after conducting a permutation test. However, when examining pseudo-F2 plants from cross B, significant QTLs with high effects were found in multiple scaffolds exceeding the LOD threshold of 6.7. The scaffolds include scaffold 10 with LOD of 11.6, scaffold 81 with LOD of 10.1, scaffold 6 with LOD of 9.5, and scaffold 14 with LOD of 8. Each QTL was accompanied by other significant QTLs nearby (Fig. **5b**, Tab. **S6**). For instance, in scaffold 10, there were three QTL peaks. The biggest peak had a LOD score of 11.6 at position 11,877,263, and two others peaks had LOD scores of 8.4 and 7.9 at positions 10,336,852 and 10,337,656, respectively. The same happened in scaffolds 81, 6, and 14 with nearby peaks. Combining the datasets from cross A and cross B increased the sample size and enhanced the statistical power for the analysis. As a result, the genome scan revealed a more robust QTL effect on scaffold 10, precisely at the same genomic location 11,877,263. The QTL exhibited a higher LOD score of 12.2, indicating a stronger and more significant association with the resistance trait (Tab. **S6**). Additional QTLs were found on scaffolds 6, and 14, but with lower effects.

**Fig. 5.**
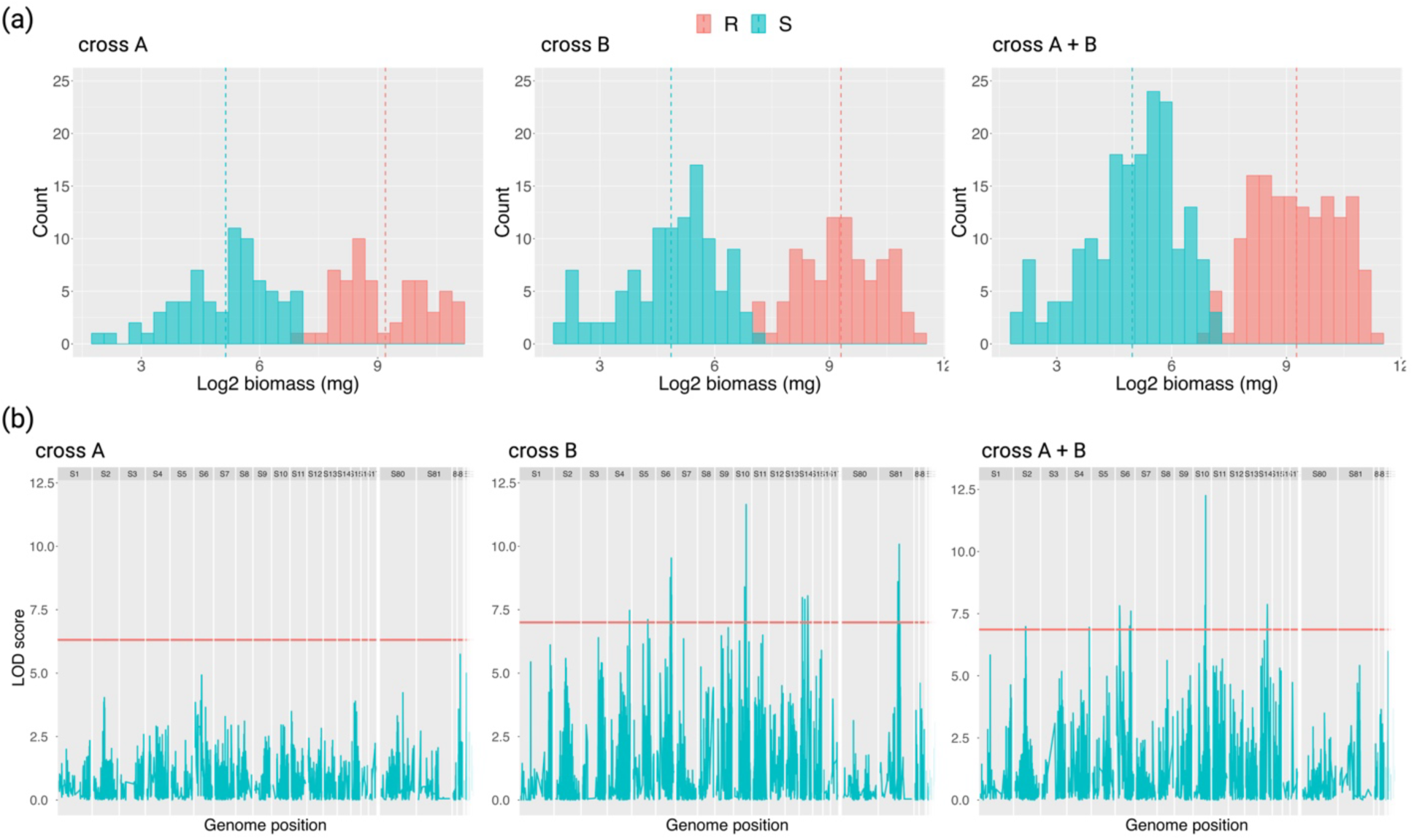
Frequency distribution and genome scan analysis. (a) Distribution of fresh biomass in plants used for QTL mapping of pseudo-F2 populations from cross A, B, and combined A+B. Blue and red colors represent sensitive and resistant pseudo-F2 plants, respectively. Dashed line indicates the average biomass for parental NES (blue) and parental NER (red). (b) Genome scan of pseudo-F2 *A. palmeri* populations from cross A, B, and combined A+B. y-axis indicates LOD score (logarithm of the odds) statistical estimate, x-axis indicates genome position on scaffolds levels. The plants were treated with 77g ha^−1^ of tembotrione, and fresh biomass was measured after 28 d.

QTL located in scaffold 10 accounted for 23% of the phenotypic variation observed in the data for cross B. Additionally, the QTLs in scaffold 81, 6, and 14 explained 20, 19, and 16% of the phenotypic variation, respectively. However, when the data from both cross A and cross B were combined, the overall phenotypic variation explained by the QTLs slightly decreased. This can be attributed to the absence of any QTL in cross A. Specifically, for the combined data, the QTL in scaffold 10 explained 15% of the phenotypic variation, while the QTLs in scaffold 6 and 14 accounted for 10% each (Tab. **S6**).

To conduct further analysis, functional annotation was performed on the combined dataset due to its larger number of genes and sample size. The QTLs found in scaffolds 10, 6, and 14 are localized on chromosomes four, eight, and fifteen of *A. palmeri*, respectively, and encompass 82, 196, and 25 genes, respectively (Supplemental Information Tab. **S6**). The QTL on scaffold 10/chromosome 4 does not contain the *CYP72A1182* gene but is located 3 Mb away. Functional annotation of genes in this QTL indicates a predominant molecular function of protein binding (GO:0005515), RNA binding (GO:0003723), and mRNA binding (GO:0003729), consistent with proteins involved in regulating other proteins or molecules through selective protein interactions (Hudson and Ortlund 2014) (Fig. **S15**). The important TFs that were identified include *DREB2A*, *WRKY* and GATA16 within the QTLs. The QTLs also include nine zinc finger proteins, four ubiquitin proteases, three F-box proteins, three ABC transporters, and three cytochrome P450 enzymes.

For each QTL, multiple markers were identified. In the case of the QTL in scaffold 10, a total of 54 markers were found, while for the QTL in scaffold 6, 94 markers were identified. However, for the QTL in scaffold 14, only a single marker was detected within a 1 Mb range upstream and downstream of the QTL region. The markers associated with scaffolds 10 and 6 displayed a consistent pattern with the evaluated phenotype, where the RR genotype exhibited higher biomasses, while the SS genotype had lower biomasses (Fig**. 6**). In contrast, the single marker found in scaffold 14 did not demonstrate the same consistency as the markers identified in the other QTLs (Fig**. 6i**).

**Fig. 6.**
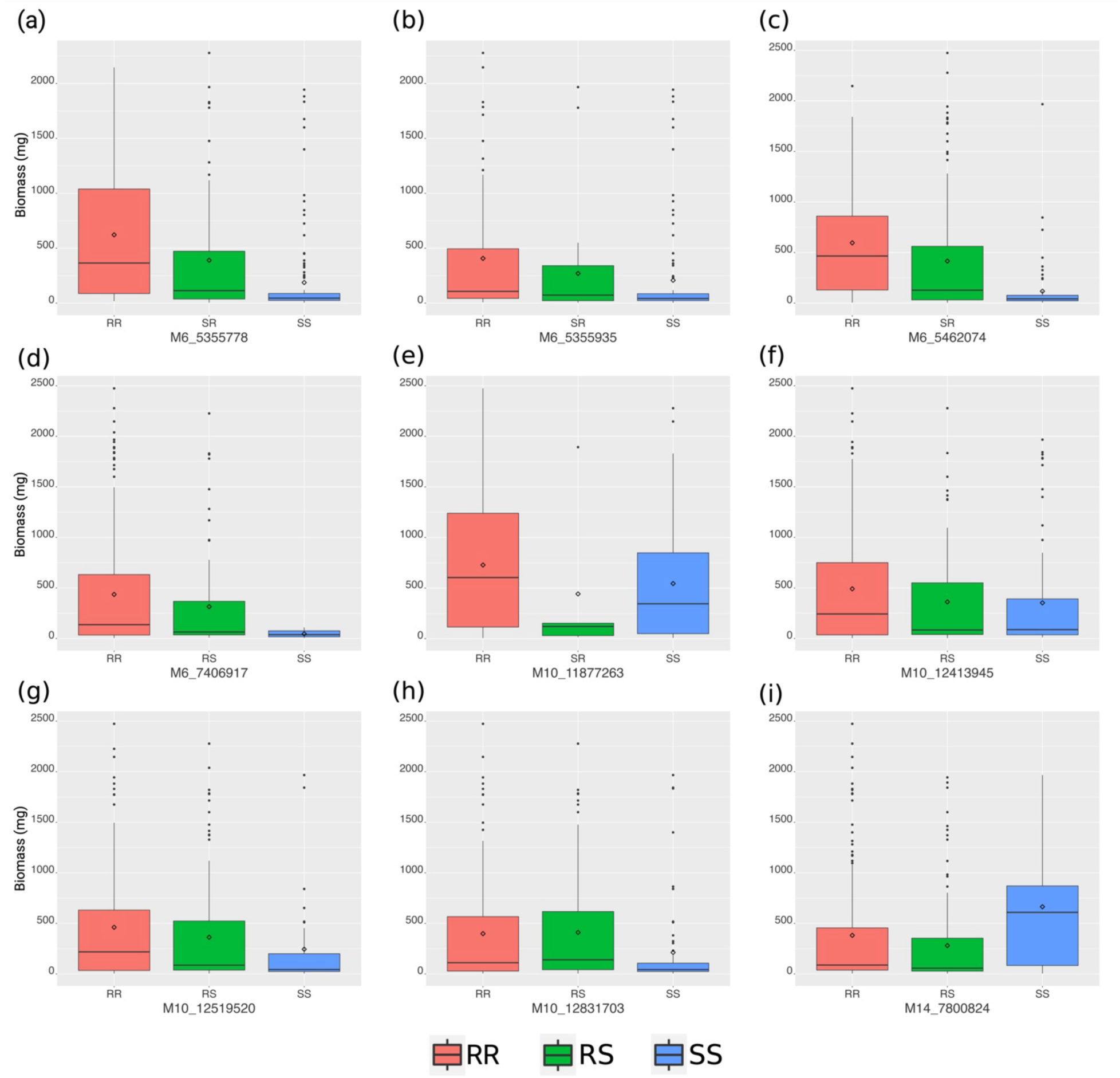
Markers and resistance phenotypes associated with QTLs for tembotrione resistance in *A. palmeri*. (a) – (d) Markers corresponding to the QTL in scaffold 6 (M6). (e) – (h) Markers corresponding to the QTL in scaffold 10 (M10). (i) Marker in scaffold 14 (M14). Boxplots represent the range of biomass values after herbicide treatment, including the maximum and minimum values, lower and upper quartiles, and the median. The diamond-shaped symbols indicate the average biomass for each genotype.

## DISCUSSION

The evolution of metabolic herbicide resistance mechanisms in weeds represents a significant challenge in weed control practices (Rigon et al. 2020). These mechanisms enable weeds to metabolize a diverse array of herbicides from various chemical families. Identifying the genes responsible for metabolic resistance and comprehending their regulation holds immense potential for developing improved herbicides and mitigating herbicide resistance. In our current investigation, four consistently up-regulated P450 genes were identified in HPPD-inhibitor resistant *A. palmeri*. These four genes did not have any polymorphisms in the coding sequences between R and S plants. Among these genes, only one, named *CYP72A1182*, deactivated the herbicide tembotrione when tested in a heterologous system. Furthermore, our study revealed that this specific gene is upregulated in other HPPD-inhibitor resistant *A. palmeri* populations collected from various fields across the US, supporting its involvement in conferring herbicide resistance in this globally significant species.

In the past, metabolic herbicide resistance in dicot weeds was relatively uncommon, with most documented cases of metabolic resistance and identified P450 genes occurring in monocots. However, recent studies have reported instances of metabolic resistance in dicots such as *A. tuberculatus* in 2017 (Kaundun et al. 2017), *A. palmeri* (Küpper et al. 2018) in 2018, and *Raphanus raphanistrum* in 2020 (Lu et al. 2019). These weed species exhibited resistance to HPPD inhibitors. Metabolic resistance to 2,4-D was identified in *A. tuberculatus* (de Figueiredo et al. 2022) . The identification of P450 genes capable of metabolizing herbicides in dicots is relatively limited. For example, *DsCYP77B34* from *Descurainia sophia*, a broadleaf weed found in Asian wheat fields, conferred resistance to ALS-, PPO-, VLCFA-, and PSII-inhibitors when expressed in *Arabidopsis thaliana* (Shen et al. 2022). Arabidopsis lines expressing *R. raphanistrum* genes *RrCYP704C1* or *RrCYP709B1* were resistant to HPPD-inhibitors such as mesotrione *(RrCYP704C1)*, tembotrione *(RrCYP709B1)*, and isoxaflutole *(RrCYP709B1)* (Lu et al. 2023). In *A. palmeri*, *CYP81CJ2* was reported as the primary gene responsible for metabolic resistance in an HPPD-inhibitor resistant population from Kansas, USA (KCTR) (Aarthy et al. 2022). The population in that study exhibited enhanced metabolism of tembotrione, and *CYP81CJ2* expression, measured by qPCR, was implicated as the likely cause of tembotrione resistance. However, our study indicates that although *CYP81CJ2* is up-regulated, it is not the causal gene for tembotrione resistance in the NER population. Instead, we have identified *CYP72A1182* as a gene capable of converting tembotrione to hydroxy-tembotrione. This finding represents the first study which mechanistically validated a gene responsible for herbicide metabolism in *A. palmeri*. Given the complex genetic structure indicated by our QTL mapping study, additional genes may be functionally involved in conferring metabolic resistance to tembotrione and this question will be the subject of future research.

*CYP81E* has been hypothesized as the gene responsible for 2,4-D metabolic resistance in *A. tuberculatus* (Giacomini et al. 2020). This specific cytochrome P450 gene co-segregated with 2,4-D resistance based on RNA-seq analysis conducted on F2 individuals. However, the gene did not co-segregate with HPPD resistance, indicating that the genetic association was specific to 2,4-D resistance. Mechanistic gene validation for function in 2,4-D metabolism was not performed (Giacomini et al. 2020). Interestingly, *CYP81E8* and *CYP81CJ2* alleles from *A. tuberculatus* and *A. palmeri*, respectively, exhibit high sequence similarity, greater than 92%. A similar behavior of *CYP81CJ2* was observed in this study, with higher expression levels in resistant plants of *A. palmeri*; however, when tested in a heterologous system, this gene did not inactivate tembotrione. Based on this finding, we hypothesized that the NER population might have resistance to 2,4-D due to up-regulation of *CYP81CJ2*. The NER population had a resistance index ranging from 2 to 2.5, but complete control was achieved when 500 a.e. g ha^−1^ of 2,4-D was applied (data not shown). This suggests that the NER population may indeed be evolving resistance to 2,4-D; however, resistance to tembotrione is not attributed to *CYP81CJ2*, but rather to *CYP72A1182*.

Previous research has highlighted *CYP72A219* as a gene involved in herbicide resistance in weeds. A study indicated that among other metabolic genes, *CYP72A219* had constitutive up-regulation in glufosinate-resistant *A. palmeri* plants, with eight times higher expression levels in resistant plants; however, the study did not perform functional gene validation (Salas-Perez et al. 2018). Furthermore, *BsCYP72A219*, along with *BsCYP81Q32* from *Beckmannia syzigachne*, were up-regulated in mesosulfuron-methyl-resistant plants (Wang et al. 2023). These genes were validated in transgenic rice, where only *CYP81Q32* conferred resistance to ALS-inhibitors. Notably, up-regulation of two genes with only one gene having validated metabolic function for herbicide resistance, resembled the observed pattern in the present study. Additionally, an alignment of *CYP81CJ2* from *A. palmeri* revealed a 92.7% identity with *CYP81Q32-like* from *Amaranthus tricolor* (XP_057543510.1), a recently sequenced *Amaranthus* species (Wang et al. 2022). This finding indicates a very close relationship between *CYP81CJ2* and *CYP81Q32* and may even represent the same gene, designated with different names. The pattern of co-segregation and up-regulation of these genes in different weed species suggests the presence of common *cis* or *trans*-elements that potentially regulate expression of multiple P450 genes in resistant plants.

Distinct promoter sequences for *CYP72A1182* were found in R pseudo-F2 plants compared to S, while no such variations were observed for *CYP81CJ2*. QTLs with high effects on the herbicide resistance trait, localized in chromosomes four, eight, and fifteen were also identified. These QTLs include three cytochrome P450 genes and ABC transporters. While these are important genes and warrant further analysis, a gene responsible for metabolizing tembotrione was identified through RNA-seq analysis and validated, suggesting that the new P450s in the QTL mapping may have a small effect for the resistance mechanism of the NER population. The key findings from the QTL are the diverse set of regulatory genes, including key transcription factors that could play a crucial role in the up-regulation of *CYP72A1182* in resistant plants. Transcription factors such as *DREB2A*, *WRKY,* and *GATA16*, along with zinc finger family found in the mapping analysis are known for their involvement in abiotic stress responses (Banerjee and Roychoudhury 2015; Seale 2020; Zhang et al. 2015). Significantly, it should be highlighted that the identified QTL on chromosome four is located two to three Mbp upstream of the *CYP72A1182* gene. This proximity might suggest that the transcription factor (TF) within the QTL region may regulate the gene, potentially influencing its expression. This regulatory effect could be particularly relevant due to the presence of a distinct promoter sequence observed in resistant plants.

Promoters of P450s have been investigated in other weed species. Nucleotide polymorphisms were observed in the promoters of *CYP81A12* and *CYP81A21* in the R alleles of multiple herbicide resistant *Echinochloa phyllopogon* plants; however, these single nucleotide polymorphisms did not exhibit a significant association with herbicide resistance (Iwakami et al. 2014). Three SNPs were identified in the promoter region of *BsCYP81Q32* in *B. syzigachne* between the R and S variants, spanning approximately 567 to 1207 base pairs upstream of the gene. These SNPs were responsible for increased expression of *GUS* in tobacco leaves transformed with the gene, suggesting the presence of a crucial sequence within that specific region (Wang et al. 2023). Additionally, the researchers identified a transcription factor, *BsTGAL6*, belonging to the bZIP family, which demonstrated efficient binding to the promoter of *BsCYP81Q32* (Wang et al. 2023). This finding suggests the involvement of a *trans*-element that regulates the expression of *BsCYP81Q32* in resistant plants.

Our study successfully identified and validated the metabolic resistant gene, *CYP72A1182*, in the tembotrione-resistant *Amaranthus palmeri* population NER. This gene demonstrated the ability to metabolize tembotrione into the main metabolite hydroxy-tembotrione. We have evidence that this gene is highly upregulated in various *A. palmeri* populations across the United States, which poses a potential threat to weed management strategies targeting this species. We also discovered important QTLs associated with the resistance trait, warranting further investigation and validation for the *CYP72A1182* regulation. Future research focusing on validating the identified transcription factors will shed light on the regulation of P450 genes and enhance our understanding of their regulation in weed species.

## ACKNOWLEDGEMENTS

This work was supported by funding from the Weed Resistance Competence Center at Bayer AG and USDA National Institute of Food and Agriculture, Hatch projects COL00783 and COL00785 to the Colorado State University Agricultural Experiment Station.

## AUTHOR CONTRIBUTIONS

CAGR, AK, AP-J, PJT, RB, FED, and TAG designed the research. CAGR, AK, CS, FP, and SS performed experiments. AK, AP-J, PJT and RB contributed experimental materials. CAGR, FP, SS, and JM analyzed the data. CAGR, FED, and TAG wrote the paper. All authors contributed to editing and approving the paper.

## DATA AVAILABILITY

The transcriptome and genome sequencing data that support the findings of this study are available at https://www.ncbi.nlm.nih.gov, with reference numbers GSE243128 and PRJNA1012341, respectively. The gene sequences are available at Genbank accession numbers OR596705-OR596711.

