## Supporting Information for "Function of Cytochrome P450 *CYP72A1182* in Metabolic Herbicide Resistance Evolution in *Amaranthus palmeri* Populations"

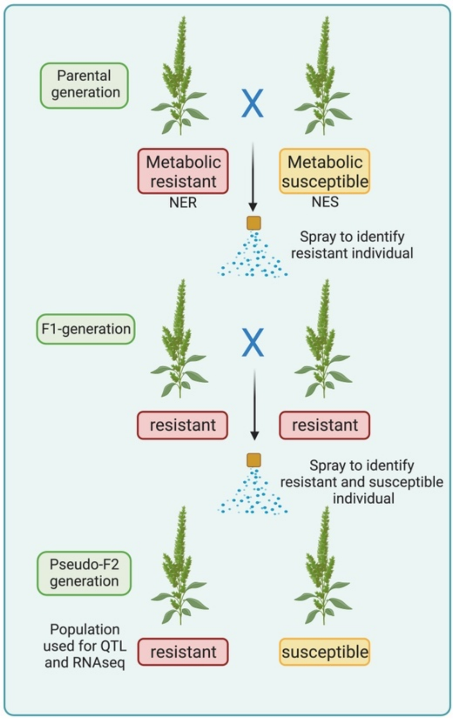


**Fig. S1.** Scheme of crosses performed to generate Pseudo-F2 generation from *Amaranthus palmeri* population HPPD-resistant (NER) x susceptible (NES).

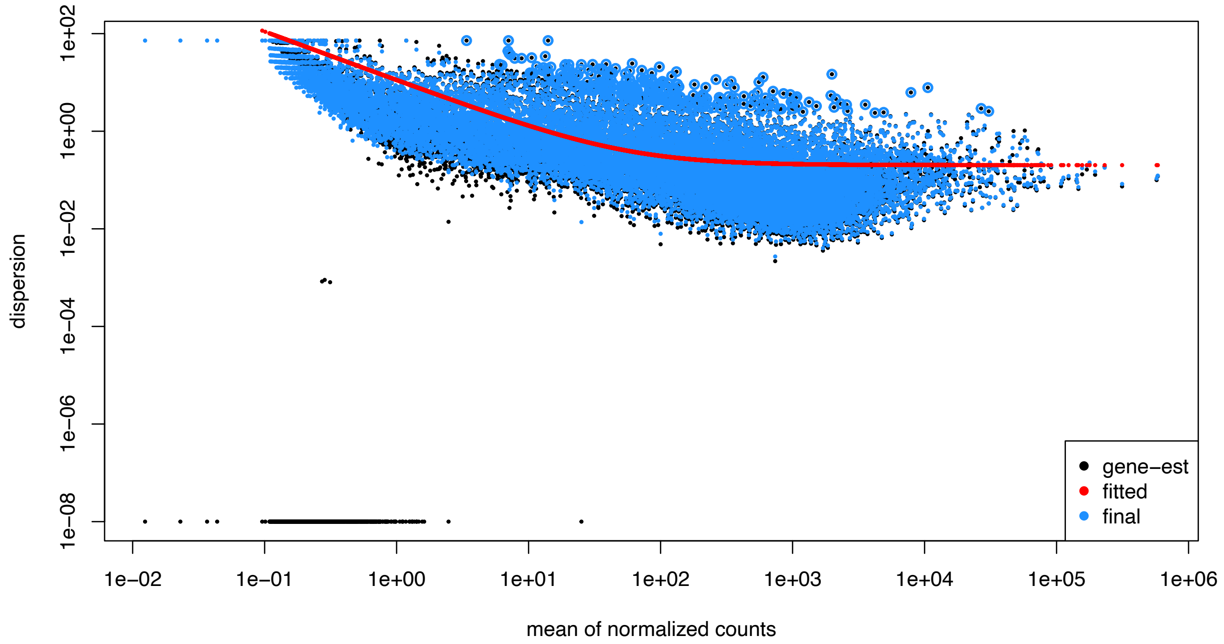


**Fig. S2.** Gene-wise dispersion estimates. The dots indicate every gene expression and its estimates toward the fitted curve. Genes with extremely high dispersion are not shrunk toward the curve.

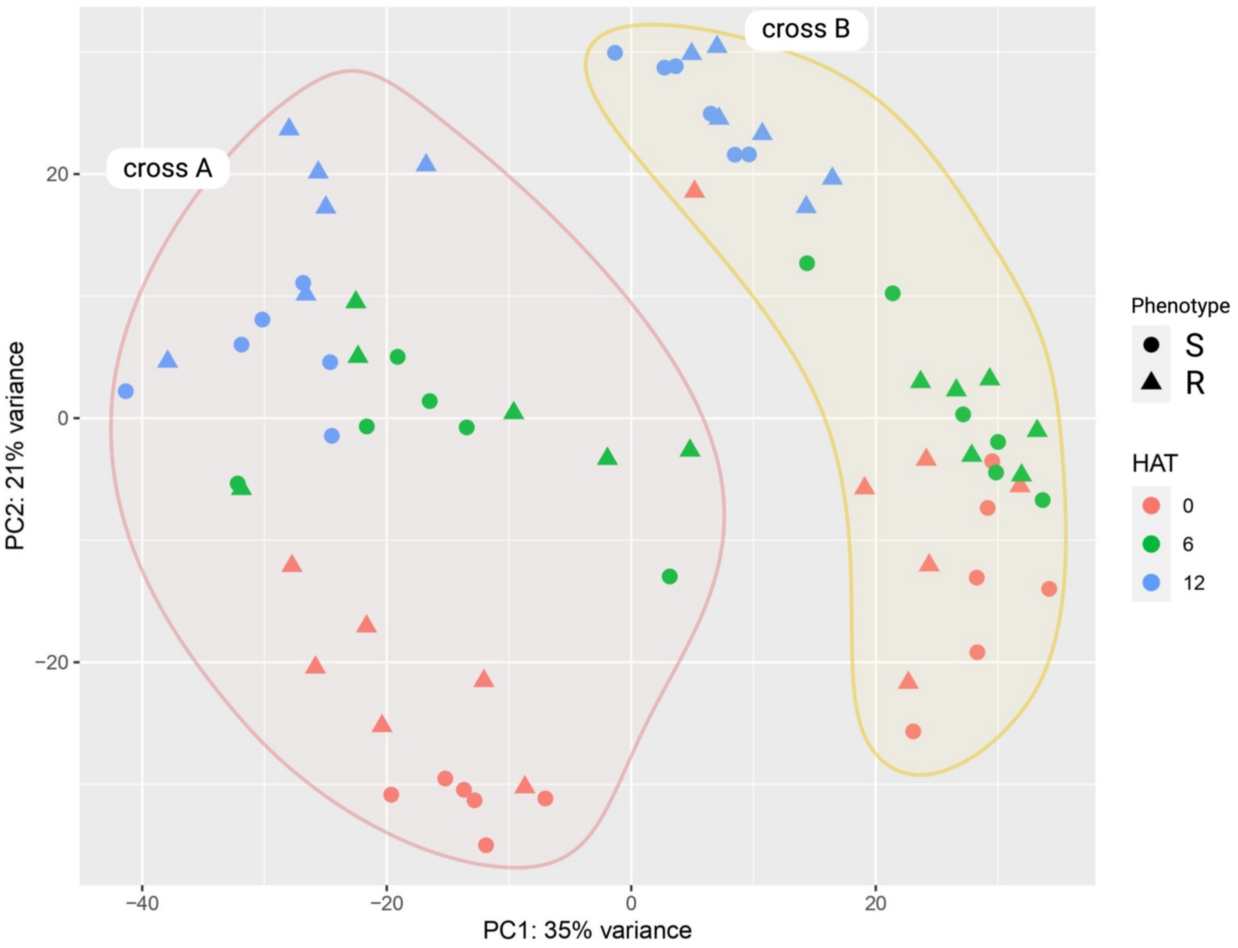


**Fig. S3.** Scatterplot of principal components 1 and 2 (35% and 21 % variance explained, respectively) for two different crosses (A and B) between sensitive (S) and HPPD-resistant (R) *Amaranthus palmeri* plants in response to tembotrione before, 6 and 12 HAT.

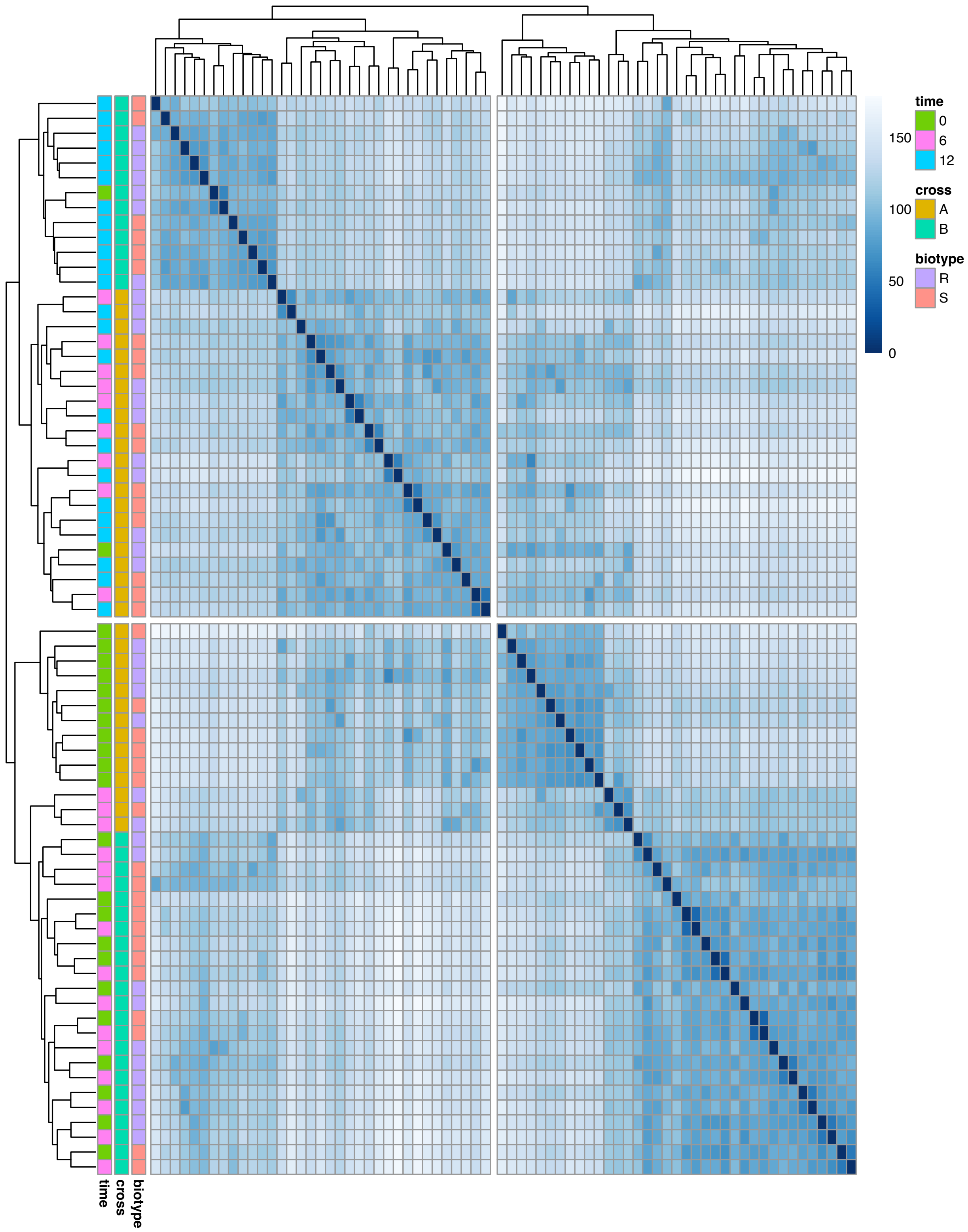


**Fig. S4.** Distance hierarchical clustering heatmap of 72 transcriptomes of *Amaranthus palmeri* generated by two crosses (A and B) between sensitive (S) and HPPD-resistant (R) in response to tembotrione application before, 6 and 12 HAT.

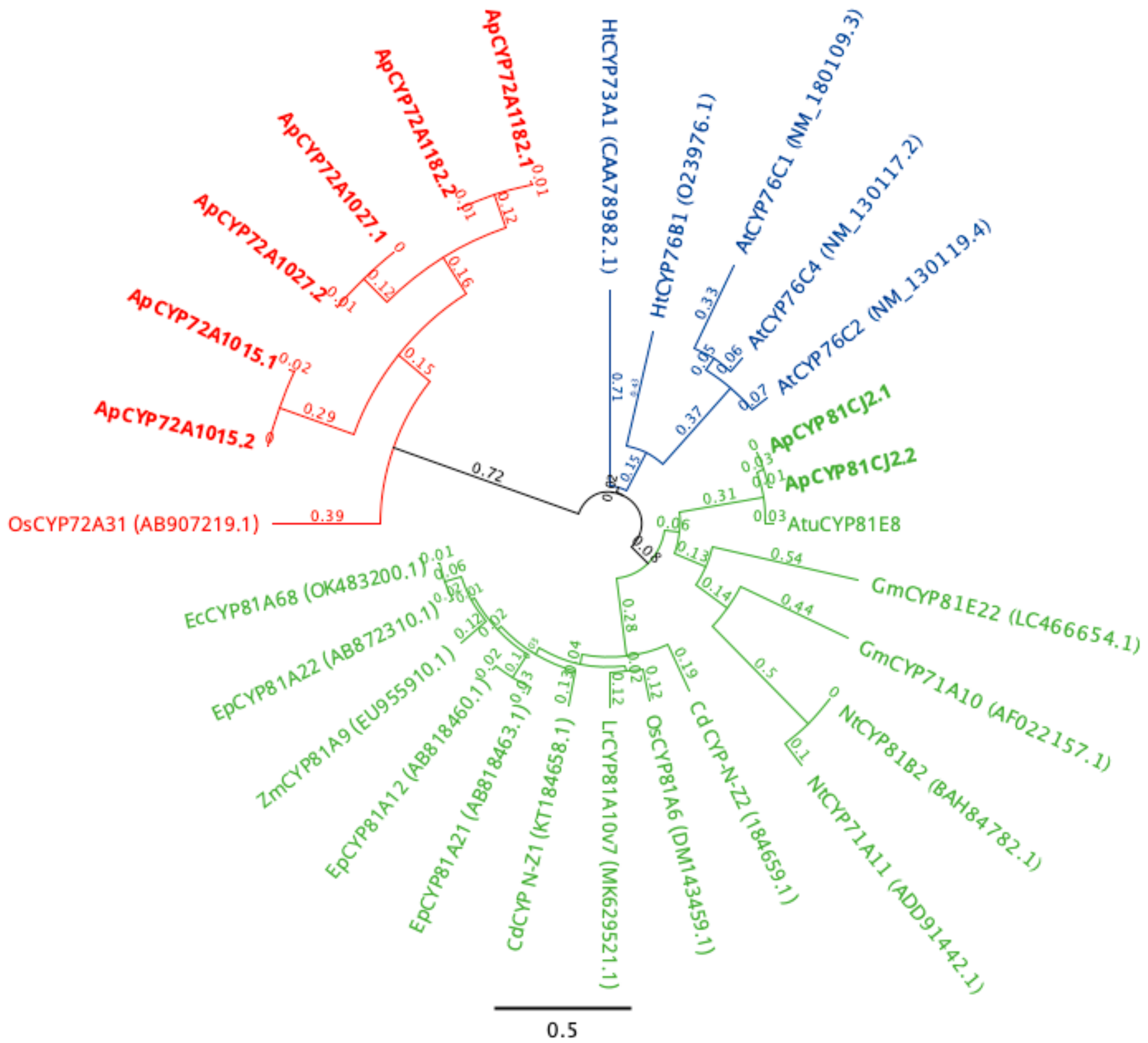


**Fig. S5.** Phylogenetic analysis of CYP72A1182, CYP72A1027, CYP72A1015 and CYP81CJ2 proteins from *Amaranthus palmeri* and other P450s that metabolize herbicides from different species. Ap – *Amaranthus palmeri*, Atu – *Amaranthus tuberculatus*, At – *Arabidopsis thaliana,* Cd – *Cynodon dactylon*, Ec – *Echinochloa crus-galli*, Ep – *Echinochloa phyllogon*, Gm – *Glycine max*, Ht – *Helianthus tuberorus*, Lr – *Lolium rigidum*, Nt – *Nicotiana tabacum*, Os – *Oryza sativa*, Zm – *Zea mays.*

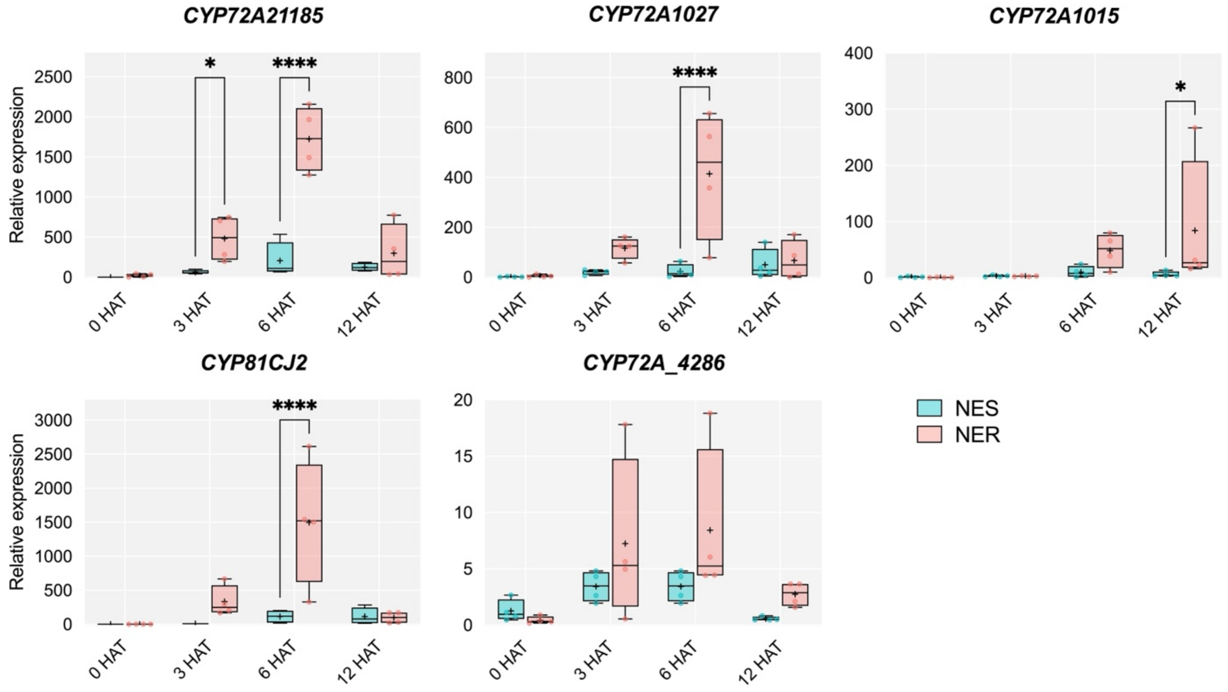


**Fig. S6.** Relative gene expression of candidate cytochrome P450 genes in parental NES and NER plants. Boxplot indicates the average, the minimum and maximum values obtained from four biological and two technical replicates. Plants samples were collected before, 3, 6 and 12 h after tembotrione application (HAT) (91 g a.i. $ha^{-1}$). The relative expression values were calculated by normalizing to the gene expression of NES untreated, using the average of the normalization gene *18S* and *Actin7*. Statistical differences at each time point were determined using the Fisher's LSD test, with asterisks indicating the significance level: * $p < 0.05$, and **** $p < 0.0001$.

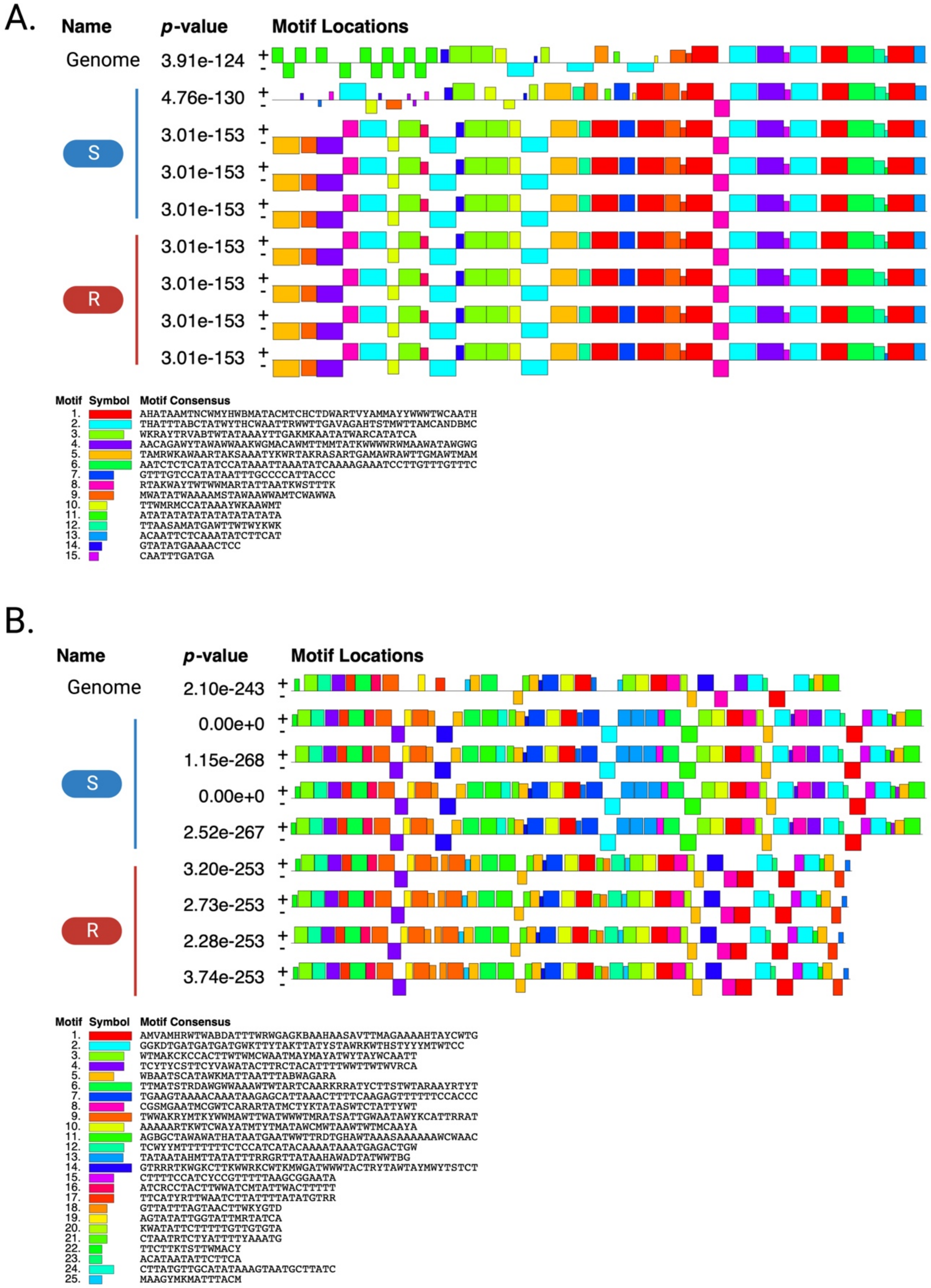


**Fig. S7.** Motifs analysis using MEME-suite tool (Bailey & Elkan, 1994) for *CYP81CJ2* and *CYP72A1182* promoter between S and R samples. Different color indicates different motif.

| Motif | Logo | Presence in promoter | Molecular Function |
|---|---|---|---|
| TCWYYMTTTTTTTCTCCATCATACAAAATAAATGAGACTGW | 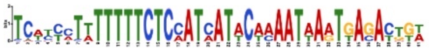 | 1x S<br>2x R | TF<br>ZI |
| TATAATAHMTTATATTTRRGRTTATAAHAWADTATWWTBG | 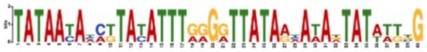 | 3x S | TF |
| GTRRRTKWGKCTTKWWRKCWTKMWGATWWWTACTRYTA WTAYMWYTSTCT | 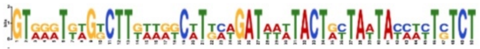 | 1x R | TF |
| GTTATTTAGTAACTTWKYGTD | 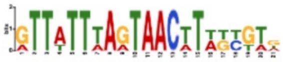 | 1x S<br>3x R | TF |
| CTAATRTCTYATTTTYAAATG | 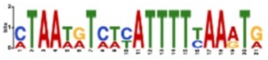 | 1x S<br>2x R | TF |
| CTTTTCCATCYCCGTTTTTAAGCGGAATA | 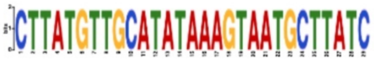 | 1x S | TF |
| MAAGYMKMATTTACM | 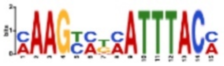 | 2x R | - |

**Fig. S8.** Motif sequences found in *Amaranthus palmeri CYP72A1182* promoter in sensitive and resistant pseudo-F2 plants. TF, transcription factor activity. ZI, zinc ion binding.

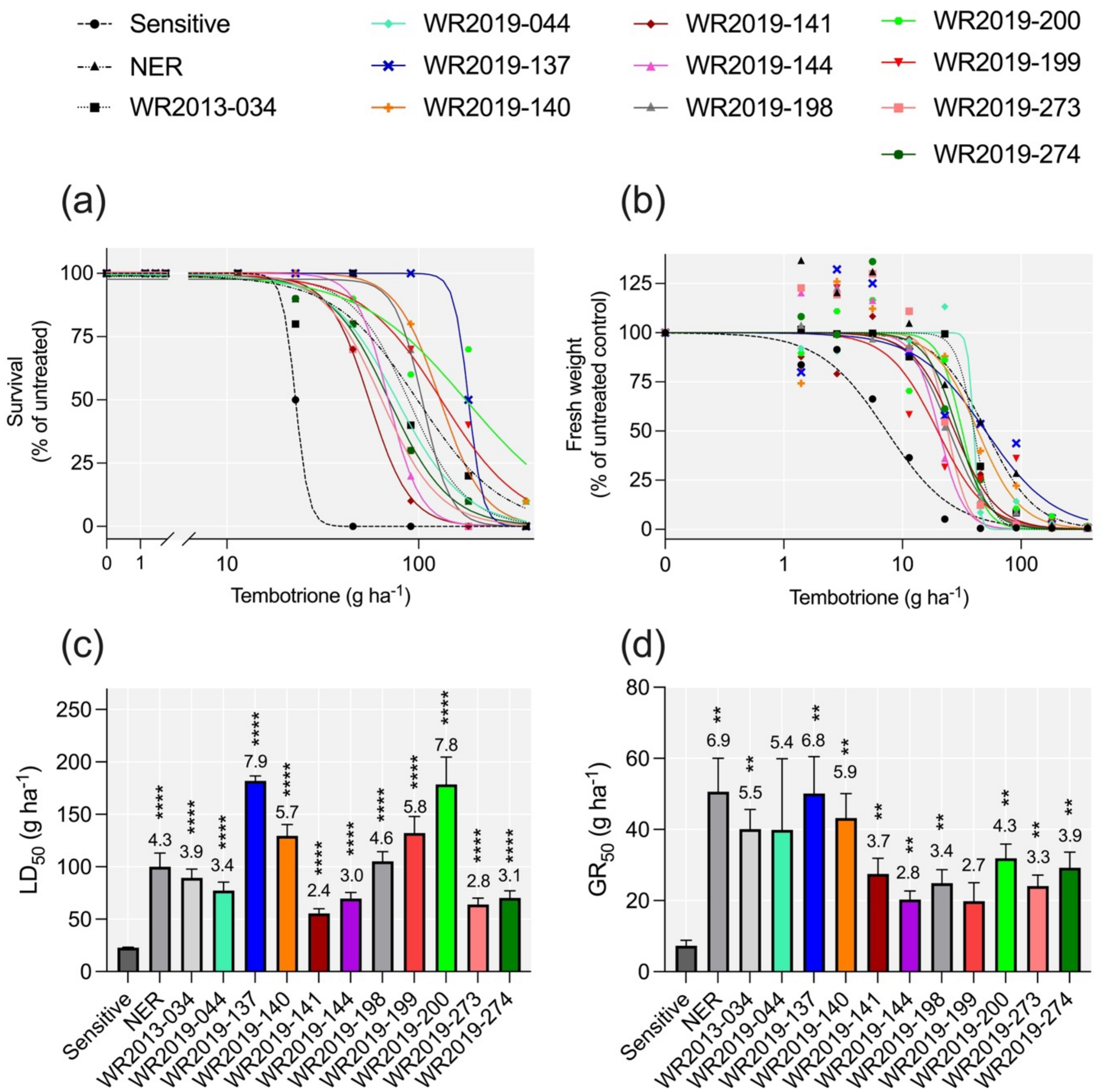


**Fig. S9.** Dose response of tembotrione in different *Amaranthus palmeri* populations. (a) Survival (% of untreated control) and (b) Fresh shoot weight (% of untreated control) of *A. palmeri* populations submitted to increasing doses of tembotrione. Field rate 91 g a.i. $ha^{-1}$. (c) Estimated $LD_{50}$ and (d) $GR_{50}$ that caused a 50 % reduction in each variable, along with the corresponding resistance index when comparing the parameter with sensitive population. Bars indicates standard error: ** <0.01, ****<0.0001.

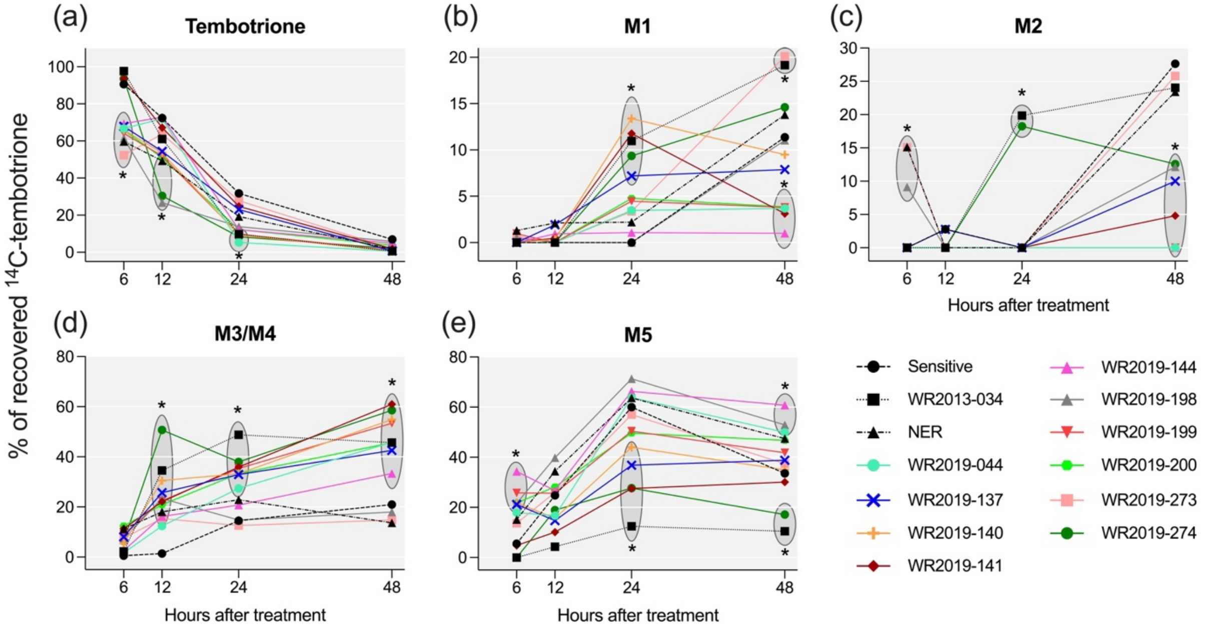


**Fig. S10.** Parental tembotrione and metabolites in HPPD resistant populations of *Amaranthus palmeri* over time after tembotrione application. Average of six plants per population and treatment. (a) Parental tembotrione, (b) Metabolite 1, Glycosylated tembotrione; (c) Metabolite 2; (d) Metabolite 3 and 4 – Hydroxylated-tembotrione. (e) Metabolite 5 – Reduced tembotrione. Asterisk indicates significant differences ($p<0.05$) between each population with sensitive by Dunnett's Test.

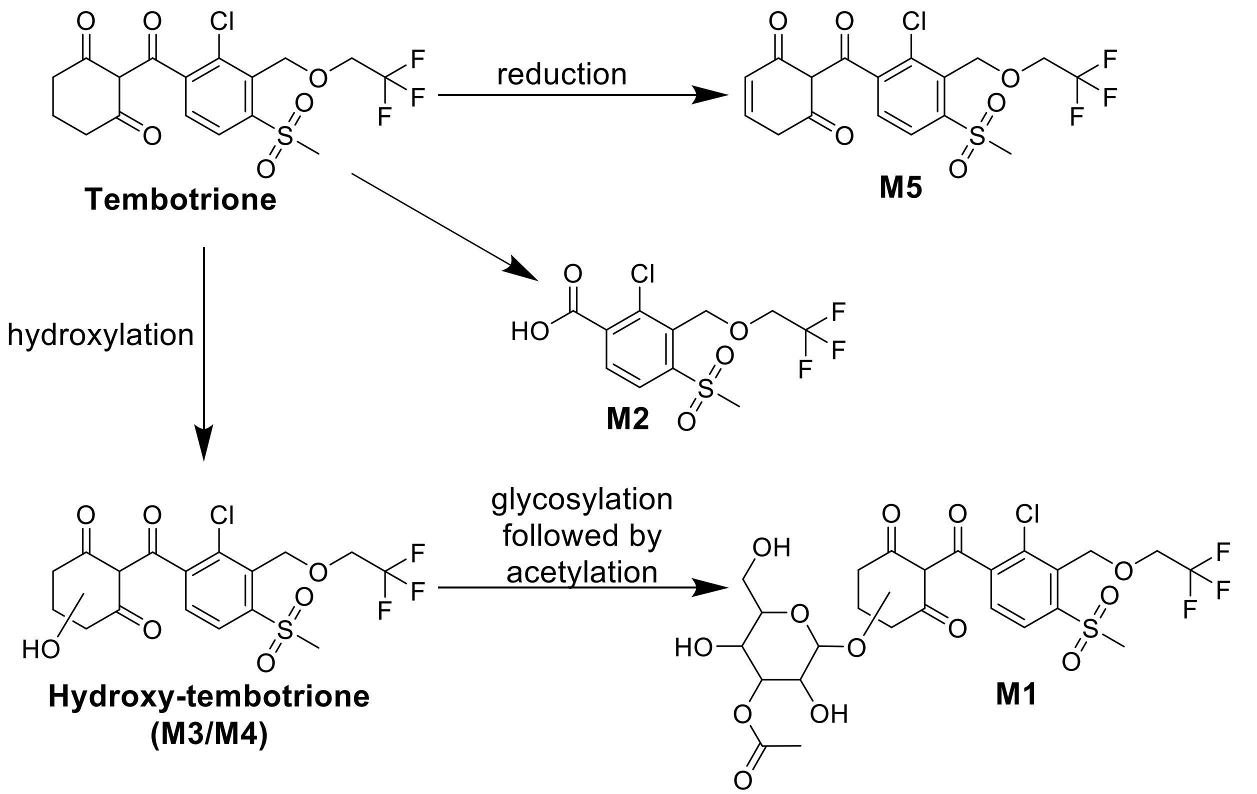


**Fig. S11.** Chemical structures of tembotrione and the main metabolites. Detoxification pathways proposed by Küpper et al. (2018). The asterisk in the tembotrione molecule marks the location of the $^{14}$C-label.

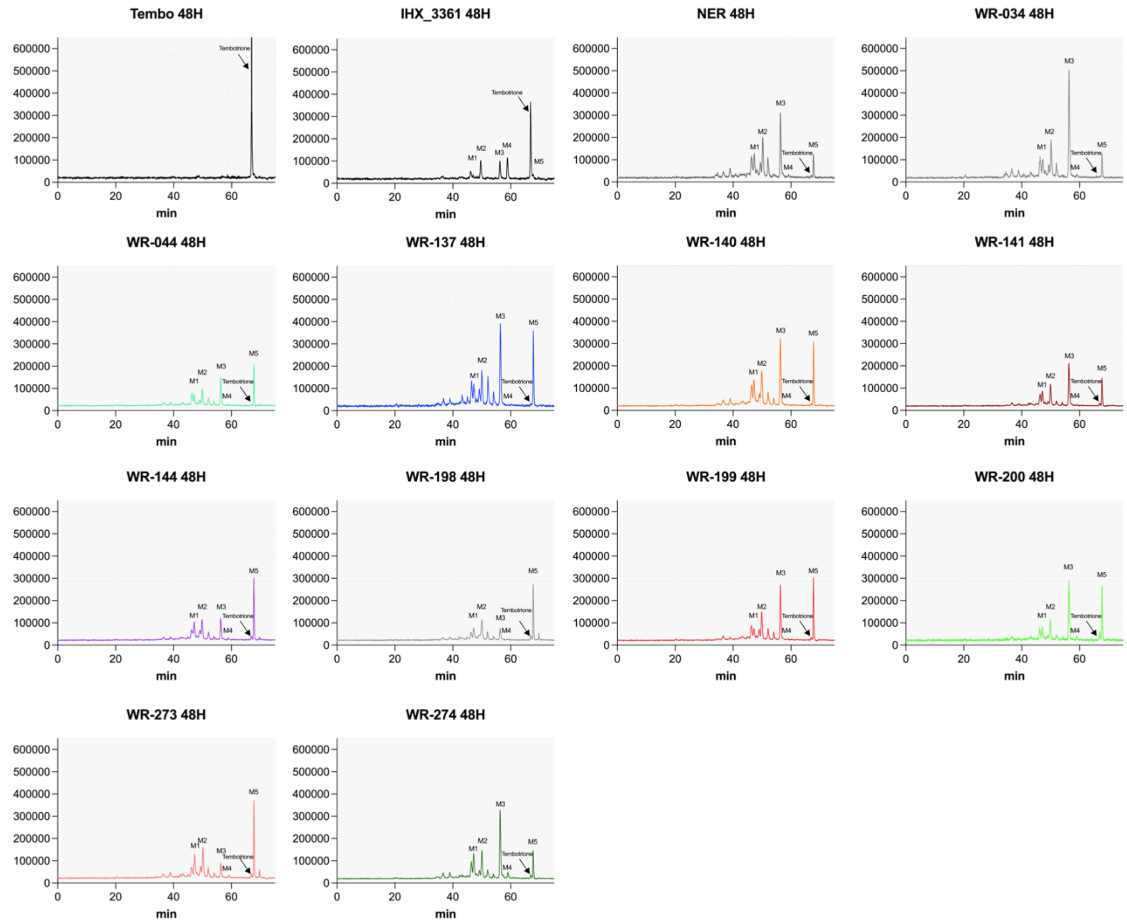


**Fig. S12.** Representative reverse-phase HPLC chromatogram for *Amaranthus palmeri* populations at 48 HAT with $^{14}C$-Tembotrione. Retention times of 47.3, 50.1, 56.3, 59.0, 66.9 and 67.7 min correspond to M1, M2, M3, M4, tembotrione and M5, respectively. IHX_3361 is herbicide sensitive control, NER and WR2013-034 are HPPD resistant control.

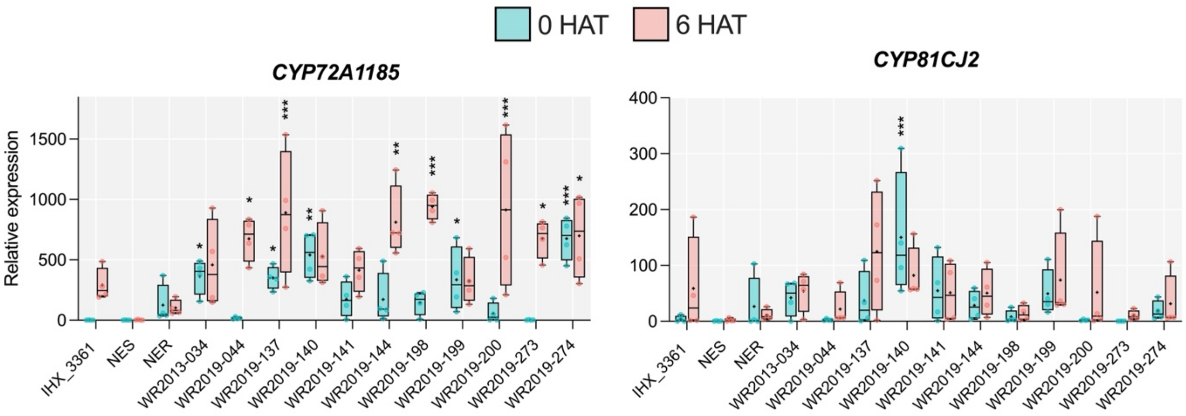


**Fig. S13.** Relative gene expression of candidate cytochrome P450 genes in different *Amaranthus palmeri* populations. Boxplot indicates the average, the minimum and maximum values of four biological and two technical replicates. Plants were collected before (0 HAT) and at 6 h after tembotrione application (HAT) (91 g a.i ha$^{-1}$). Relative expression was calculated using the average of the normalization gene *18S* and normalized relative to the gene expression in the sensitive biotype untreated. IHX_3361 and NES are herbicide sensitive control, NER and WR2013-034 are HPPD resistant control. The asterisk indicates the statistical difference between each population with sensitive for each time point by the Fisher's LSD test, * <0.05, ** <0.01, *** <0.001.

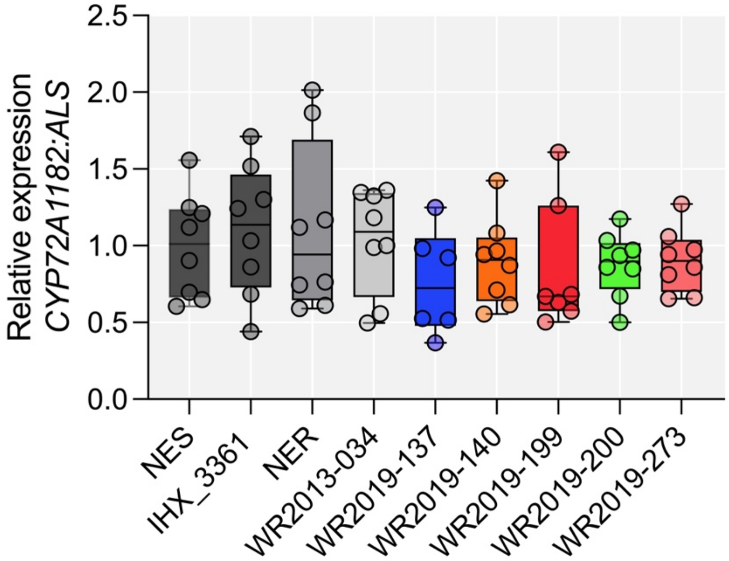


**Fig. S14.** *CYP72A1182* copy number relative to *ALS* in different populations of *Amaranthus palmeri*. NES and IHX_3361 are herbicide sensitive control, NER and WR2013-034 are HPPD resistant control. Circle indicates biological replicate.

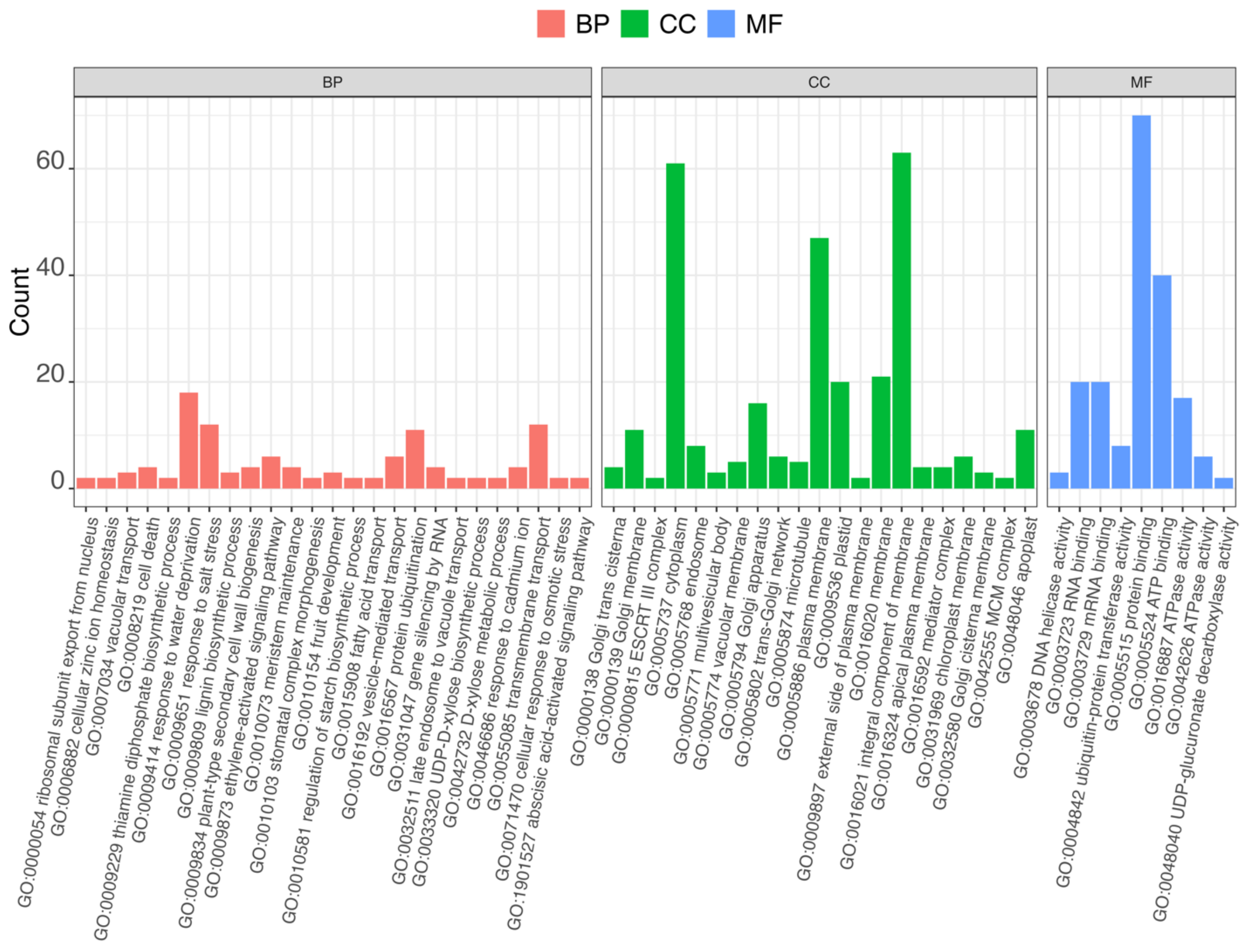


**Fig. S15.** Functional enrichment of genes from QTLs of scaffold 10, scaffold 6 and scaffold 14. Detailed information of functional annotations of all the genes based on GO classification, including biological process (BP), cellular component (CC) and molecular function (MF).

**Tab. S1.** Gene stability assessed by NormFinder algorithm.

| Gene name | | Stability value |
|---|---|---|
| *18S* | | 0.110 |
| *ACT* | | 0.193 |
| *TUB* | | 0.231 |
| Intergroup variation<br>Group identifier | NES | NER |
| *18S* | 0.068 | -0.068 |
| *ACT* | -0.149 | 0.149 |
| *TUB* | 0.080 | -0.080 |
| Intragroup variation<br>Group identifier | NES | NER |
| *18S* | 0.515 | 0.128 |
| *ACT* | 1.048 | 0.746 |
| *TUB* | 1.147 | 1.427 |
| Best gene | | 18S |
| Stability value | | 0.110 |
| Best combination of two genes | | 18S and ACT |
| Stability value for best combination of two genes | | 0.112 |

NES – Nebraska susceptible, NER – Nebraska HPPD-resistant.

**Tab. S2.** Primer sequences used for different experiments and their characteristics.

| Experiment | gene | sequence 5' to 3' | size (bp) |
|---|---|---|---|
| Gene validation | qPCR_actin_F | ggctgatgcagaggagattc | 149 |
| | qPCR_actin_R | gtcccataccaaccatgacacc | |
| | 18S rRNA_F | caaccataaacgatgccgacc | 113 |
| | 18S rRNA_R | cagccttgcgaccatactcc | |
| | TUB_F | cattatactgaaggtgctgaac | 211 |
| | TUB_R | ctttcggagatgggaaaaca | |
| | ALSF2 (Gaines *et al.*, 2010) | gctgctgaaggctacgct | 118 |
| | ALSR2 (Gaines *et al.*, 2010) | gcg ggactgagtcaagaagtg | |
| | CYP72A1182_F | gccaaaaaggttagaaaaatgc | 130 |
| | CYP72A1182_R | taggttgggcattgggttta | |
| | CYP72A1027_F | aaagcatcaaaattggcaac | 213 |
| | CYP72A1027_R | actacgacacctgctggaag | |
| | CYP72A219_4286_F | tcgaaagccaaaattgaacc | 158 |
| | CYP72A219_4286_R | ccttaacactggtgccgaat | |
| | CYP72A1015_F | gccggtgtgcaagttaaaat | 232 |
| | CYP72A1015_R | atgaaaagtgcggcaaaatc | |
| | CYP81CJ2_1F | gacaatgatcgcaccaacac | 131 |
| | CYP81CJ2_1R | cagggaacaatgtggctctt | |
| Promoter amplification | CYP81CJ2_F | ggacaaaggagtgaaatcatcg | 2131 |
| | CYP81CJ2_R | catgcggaggaaactaccac | |
| | CYP72A1182_Pro_F | gggtcactgtcttttgatttaggg | 1951 |
| | CYP72 A1182_Pro_R | tggtcatccaagctctttca | |

**Table S3.** Constitutively differentially expressed genes between tembotrione resistant and susceptible *Amaranthus palmeri*. Adjusted *P*-value <0.01.

| gene ID | log2FC | lfcSE | Gene | Molecular function |
|---|---|---|---|---|
| **MAKER_16168** | 3.57 | 0.43 | Glutathione S-transferase U19 (GSTU9) | glutathione transferase activity |
| **MAKER_12558** | 5.83 | 0.92 | Disease resistance protein At4g27190-like (LOC104598357) | ADP binding |
| **MAKER_29886** | 4.20 | 0.57 | NADPH--cytochrome P450 reductase (CYP72A219-like) | monooxygenase activity |
| **MAKER_05008** | 3.37 | 0.43 | Scopoletin glucosyltransferase (TOGT1) | scopoletin glucosyltransferase activity |
| **MAKER_08144** | 3.19 | 0.40 | glycosyltransferase | - |
| **MAKER_06269** | 22.60 | 3.43 | Protein of unknown function | - |
| **MAKER_25717** | 3.56 | 0.52 | NADPH--cytochrome P450 reductase (CYP72A219-like) | monooxygenase activity |
| **MAKER_08142** | 3.71 | 0.56 | Protein of unknown function | - |
| **MAKER_21351** | 2.47 | 0.33 | Crocetin glucosyltransferase (GLT2) | glucosyltransferase activity |
| **MAKER_20679** | 1.90 | 0.25 | Detoxification 27 (DTX27) | transmembrane transporter activity |
| **MAKER_03299** | 4.50 | 0.85 | MADS-box transcription factor 23-like | DNA-binding transcription factor activity |
| **MAKER_31185** | 21.49 | 4.09 | Protein of unknown function | - |
| **MAKER_16324** | 2.94 | 0.48 | Protein of unknown function | - |
| **MAKER_04610** | 2.20 | 0.33 | Enhanced pseudomonas susceptibility 1 (EPS1) | acyltransferase activity |
| **MAKER_28341** | 3.02 | 0.51 | Vacuolar-processing enzyme beta-isozyme (bVPE) | cysteine-type endopeptidase activity |
| **MAKER_20648** | 2.50 | 0.40 | Betanidin 6-O-glucosyltransferase | glucosyltransferase activity |
| **MAKER_10107** | 2.64 | 0.44 | Cytochrome P450 81E8 (CYP81E8) | monooxygenase activity |
| **MAKER_07960** | 2.38 | 0.39 | Glycosyltransferase GTB type superfamily protein | transferase activity |
| **MAKER_08145** | 2.69 | 0.45 | UDP-glycosyltransferase 71B2-like | UDP-glycosyltransferase activirty |
| **MAKER_04612** | 2.03 | 0.31 | acetyl-transferase-like | transferase activity |
| **MAKER_32649** | 4.15 | 0.89 | MADS-box transcription factor 27 (MADS27) | DNA-binding transcription factor activity |
| **MAKER_20644** | 2.93 | 0.54 | UDP-glycosyltransferase 71A15 | transferase activity |
| **MAKER_21756** | 3.01 | 0.57 | glutathione S-transferase (GST) | transferase activity |
| **MAKER_04611** | 2.00 | 0.34 | BAHD acyltransferase DCR-like protein | transferase activity |
| **MAKER_15874** | 3.54 | 0.80 | Protein of unknown function | - |
| **MAKER_06150** | 2.06 | 0.36 | Glutathione S-transferase U19 (GSTU9) | glutathione transferase activity |
| **MAKER_25718** | 2.43 | 0.47 | NADPH--cytochrome P450 reductase (CYP72A219-like) | monooxygenase activity |
| **MAKER_24005** | 2.16 | 0.42 | NADPH:quinone oxidoreductase | NADPH dehydrogenase activity |
| **MAKER_16167** | 2.16 | 0.44 | Glutathione S-transferase (GST) | glutathione transferase activity |
| **MAKER_34451** | 3.15 | 0.95 | Protein of unknown function | - |
| **MAKER_17494** | 3.20 | 1.05 | Protein of unknown function | - |
| **MAKER_12474** | 3.27 | 1.90 | RING-H2 finger protein ATL3 | metal ion binding |
| **MAKER_12523** | 1.55 | 0.29 | Aldehyde dehydrogenase family 7 member B4 (ALDH7B4) | catalytic activity |

| | | | | |
|---|---|---|---|---|
| **MAKER_12487** | 3.29 | 1.53 | Protein of unknown function | |
| **MAKER_12408** | 1.81 | 0.37 | Beta-amyrin synthase 1 | intramolecular transferase activity |
| **MAKER_22184** | 2.00 | 0.46 | Calcium-binding protein (CP1) | binding |
| **MAKER_27594** | 2.61 | 0.78 | Lateral Organ Boundaries (LOB) | DNA binding |
| **MAKER_11763** | -2.55 | 0.4656 | Abscisic-aldehyde oxidase (AAO3) | aldehyde oxireductase |
| **MAKER_11325** | -1.49 | 5.1123 | Protein of unknown function | - |
| **MAKER_29818** | -3.04 | 1.335 | Abscisic acid receptor (PYL4) | abscisic acid binding |

**Table S4.** Logistic equation parameters and the resistance index (RI) of survival (% of untreated control) for different populations of *Amaranthus palmeri* subjected to different doses of tembotrione.

| populations | b | d | $LD_{50}$ dose (g $ha^{-1}$) | lower CI (95%) | upper CI (95%) | SE | RI | *p*-value |
|---|---|---|---|---|---|---|---|---|
| Sensitive | 12.6 | 100.0 | 22.8 | 21.8 | 23.8 | 0.5 | - | |
| NER | 1.9 | 98.8 | 99.9 | 74.5 | 125.5 | 12.9 | 4.3 | <0.001 |
| WR2013-034 | 2.9 | 98.5 | 89.4 | 73.1 | 105.8 | 8.3 | 3.9 | <0.001 |
| W2019-044 | 2.5 | 99.5 | 77.3 | 61.6 | 93.0 | 8.0 | 3.4 | <0.001 |
| W2019-137 | 11.6 | 100.0 | 181.9 | 172.8 | 191.1 | 4.6 | 7.9 | <0.001 |
| W2019-140 | 3.8 | 100.3 | 129.4 | 108.2 | 150.6 | 10.8 | 5.7 | <0.001 |
| W2019-141 | 4.1 | 98.9 | 55.4 | 46.8 | 64.5 | 4.4 | 2.4 | <0.001 |
| W2019-144 | 5.2 | 100.0 | 69.6 | 57.8 | 81.3 | 5.9 | 3.0 | <0.001 |
| W2019-198 | 6.2 | 97.3 | 105.2 | 87.2 | 123.24 | 9.1 | 4.6 | <0.001 |
| W2019-199 | 2.1 | 99.5 | 132.1 | 100.4 | 163.2 | 15.8 | 5.8 | <0.001 |
| W2019-200 | 1.5 | 99.0 | 178.3 | 127.6 | 229.6 | 25.9 | 7.8 | <0.001 |
| W2019-273 | 2.8 | 100.6 | 63.8 | 51.9 | 75.9 | 6.1 | 2.8 | <0.001 |
| W2019-274 | 2.8 | 99.5 | 70.3 | 57.1 | 83.6 | 6.7 | 3.1 | <0.001 |

b: slope; d: upper limit; $LD_{50}$: herbicide dose that causes a 50% reduction in survival, CI: confidence interval of the parameter $LD_{50}$ ($\alpha = 0.05$); and RI: resistance index = $LD_{50}$ ratio between respective population and sensitive.

**Table S5.** Logistic equation parameters and the resistance index (RI) of shoot fresh weight (of untreated control) for different populations of *Amaranthus palmeri* subjected to different doses of tembotrione.

| Populations | b | d | GR50 dose (g ha$^{-1}$) | lower CI (95%) | upper CI (95%) | SE | RI | *p*-value |
|---|---|---|---|---|---|---|---|---|
| Sensitive | 1.5 | 100 | 7.3 | 4.3 | 10.4 | 1.5 | - | - |
| NER | 2.1 | 100 | 50.6 | 32.2 | 69.0 | 9.3 | 6.9 | <0.01 |
| WR2013-034 | 5.7 | 100 | 40.1 | 29.2 | 51.1 | 5.5 | 5.5 | <0.01 |
| W2019-044 | 17.9 | 100 | 39.9 | 0.65 | 79.1 | 19.9 | 5.4 | 0.1 |
| W2019-137 | 1.5 | 100 | 50.2 | 29.6 | 70.6 | 10.4 | 6.8 | <0.01 |
| W2019-140 | 2.4 | 100 | 43.2 | 29.6 | 70.6 | 6.9 | 5.9 | <0.01 |
| W2019-141 | 2.5 | 100 | 27.5 | 18.9 | 36.1 | 4.4 | 3.7 | <0.01 |
| W2019-144 | 4.1 | 100 | 20.3 | 15.6 | 25.0 | 2.4 | 2.8 | <0.01 |
| W2019-198 | 2.6 | 100 | 24.9 | 17.4 | 32.4 | 3.8 | 3.4 | <0.01 |
| W2019-199 | 2.1 | 100 | 19.8 | 9.5 | 30.0 | 5.2 | 2.7 | 0.06 |
| W2019-200 | 4.5 | 100 | 31.9 | 23.9 | 39.8 | 4.0 | 4.3 | <0.01 |
| W2019-273 | 5.1 | 100 | 24.1 | 17.8 | 30.2 | 3.1 | 3.3 | <0.01 |
| W2019-274 | 3.3 | 100 | 29.2 | 20.6 | 37.8 | 4.4 | 3.9 | <0.01 |

b: slope; d: upper limit; $GR_{50}$: herbicide dose that causes a 50% reduction in shoot fresh weight, CI: confidence interval of the parameter $GR_{50}$ ($\alpha$ = 0.05); and RI: resistance index = $GR_{50}$ ratio between respective population and sensitive.

**Table S6.** Quantitative trait loci found in Pseudo-F2 *Amaranthus palmeri* in cross B and combined cross A + B for the trait of HPPD resistance. Position are based on available genome draft of *A. palmeri* (Montgomery *et al.*, 2020).

| QTL | LOD score | Position | Interval | | Chromosome | Annotated genes | PVE |
|---|---|---|---|---|---|---|---|
| **Cross B** | | | | | | | |
| Scaffold_10 | 11.6 | 11877263 | 11865155 | 11890395 | 4 | 77 | 23 % |
| | 8.4 | 10336852 | 10224431 | 10336879 | | | |
| | 7.9 | 10337656 | 10336879 | 10482794 | | | |
| Scaffold_81 | 10.1 | 19632215 | 18232833 | 20375576 | 2 | 78 | 20 % |
| | 8.6 | 18232833 | 18186955 | 18238951 | | | |
| Scaffold_6 | 9.5 | 16851238 | 15842545 | 16851238 | 8 | 7 | 19 % |
| | 8.7 | 15843735 | 15752153 | 15907859 | | | |
| Scaffold_14 | 8.0 | 7800824 | 2267056 | 7800824 | 15 | 50 | 16 % |
| | 7.9 | 4777664 | 4754252 | 4917017 | | | |
| | 7.9 | 2267056 | 2226097 | 2271946 | | | |
| **combined cross A + B** | | | | | | | |
| Scaffold_10 | 12.25 | 11877263 | 10877263 | 12877263 | 4 | 105 | 15 % |
| | 7.84 | 11332443 | | | | | |
| Scaffold_6 | 7.81 | 5462074 | 4962074 | 5962074 | 8 | 32 | 10 % |
| | 7.60 | 16851238 | 15150809 | 17351238 | | 164 | |
| | 7.00 | 15650809 | | | | | |
| | 6.68 | 15617367 | | | | | |
| Scaffold_14 | 7.87 | 7800824 | 7300824 | 8300824 | 15 | 25 | 10 % |

PVE – phenotype variation explained.

**Methods S1** Plant material for QTL mapping and RNAseq experiment

For the crosses, *Amaranthus palmeri* seeds were sown on 0.7% agar medium (Sigma-Aldrich), placed in a refrigerator at 4 °C for seven d and then germinated on a germination bench at room temperature with 16/8h day/night cycle. Germinated seedlings were transplanted into commercial potting soil (Professional Growing Mix, Sun Gro Horticulture) in 5x5 cm inserts and maintained in the greenhouse at 24±2°C temperatures and 15/9 h day/night photoperiods supplemented with metal-halide lamps (400 µmol $m^{-2}$ $s^{-1}$). Plants were watered daily. A pseudo-F2 generation was generated by first spraying parental NER individuals at 7-10 cm height with a field rate of 91 a.i. $ha^{-1}$ tembotrione (Laudis, Bayer CropScience) and 1% v/v methylated seed oil (MSO). Herbicide applications were made using an overhead track sprayer (DeVries Manufacturing) equipped with a flat-fan nozzle tip (TeeJet 8002EVS, Spraying System) calibrated to deliver 187 L $ha^{-1}$ of spray solution at 172 kPa. Surviving NER were transplanted into 22.5 cm diameter pots and individually crossed with another NES individual using pollination bags. Five crosses with NERmale × NESfemale and five crosses with NERfemale × NESmale were performed and grown to seed. The resulting F1 generations were grown out and sprayed again under the same conditions described above. Two F1 individuals from each parental cross that survived the application were crossed with each other. Seeds from the resulting Pseudo-F2 generation were subsequently used for the QTL mapping or RNA-Seq experiment. Schemes of the crosses are available in **Fig. S1**.

For the RNA-Seq experiment, plants were grown from the Pseudo-F2 generations generated from two parental NERmale x NESfemale crosses (cross A and B, respectively). Seeds were sown on 0.7% agar medium (Sigma-Aldrich), placed in a refrigerator at 4 °C for seven d and then germinated on a germination bench at room temperature with 16/8h of day/night cycle. About 150 seedlings from each Pseudo-F2 cross were transplanted into 4x4 cm inserts each and maintained in a growth chamber at 25/22°C day/night temperatures, 70% relative humidity, and 16h photoperiod with 700 µmol m-2 s-1 provided by incandescent and fluorescent bulbs. At 4-5 cm height and 4-5 leaf stage, the first fully expanded leaf from the apical meristem of each plant was cut and immediately frozen at -80°C for timepoint 0. The plants were then sprayed at 77 g a.i. $ha^{-1}$ (85% of the field rate) with 1% v/v MSO as described above to ensure resistant plants would be able to survive despite removing leaves for testing. Six HAT the second expanded leaf and twelve HAT the third expanded leaf were cut and immediately frozen at -80 °C, respectively. Visual damage and survival data were recorded after treatment for 21 d to phenotype resistant and susceptible individuals. From each pseudo-F2 cross only the six visually most resistant and the six visually most susceptible individuals were used for RNA-Sequencing (Table X).

Total RNA was extracted from frozen ground tissue using the Direct-zol™ RNA MiniPrep Plus (Zymo Research) which includes DNAse treatment. Yield and purity were measured with a NanoDrop 2000 spectrophotometer (Thermo Scientific) and RNA integrity (RIN) was measured on an Agilent 2200 Bio TapeStation system (Agilent Technologies) using Agilent High Sensitivity RNA ScreenTape. RNASeq library preparation was performed with the TruSeq stranded mRNA library prep kit (Illumina) preparing for 150 nucleotide paired-end sequencing. The 108 libraries were run on an Illumina HiSeq 4000 platform on a total of 16 lanes (2 flow cells) with seven libraries per lane, yielding 5.2 billion paired-end reads. Individual library yields were 48.2 million on average and ranged from 35.0 to 71.5 million paired-end reads. On average over 93% of sequenced nucleotides met a quality score of 30 (Q30 Phred score).

**Methods S2.** Yeast transformation

The strains WAT11 and WAT21 of *Saccharomyces cerevisiae* expressing *Arabidopsis* genes for cytochrome P450 reductase 1 and 2 (Urban *et al.*, 1997), respectively, were used as a heterologous system to test the hypothesis that the candidate P450 genes could metabolize tembotrione. Yeast cells were grown in glucose SC (-ura) agar plates. Single cells were inoculated in YPD(A) medium and transformed using a modified lithium acetate procedure (Gietz *et al.*, 1992). Transformed yeast cells were selected by glucose SC (-ura) agar plates. Yeast colony PCR to confirm the gene insertion was performed using high-fidelity PrimerStar HS DNA polymerase by heating the master mix at 94°C for 4 min before the PCR protocol.

**Methods S3.** Promoter amplification

Reverse primers were designed on the conserved sequence of the first exon for each gene. PCR was performed using PrimeSTAR® HS DNA Polymerase kit (Takara) consisting of 10 µL 5× PrimSTAR buffer ($Mg2+$ Plus), 4 µL dNTP mix (2.5 mM each), 1.5 µL of each primer (forward and reverse), 0.5 µL PrimeSTAR HS DNA Polymerase, 1 µL of DNA (50 ng) and sterile water up to 50 µL. PCR cycling conditions were initial denaturation at 98°C for 30 s, followed by 40 cycles of denaturation at 98°C for 10 s, annealing at 60°C for 15 s, and extension at 72°C for 2.5 min. PCR product was run in 1% agarose gel electrophoresis for 30 min and amplicons were sent for sequencing using the long-read sequencing technology Oxford Nanopore Technology (ONT) by SNPsaurus LLC (https://www.plasmidsaurus.com). Sequence results were confirmed by blasting the reads to the *A. palmeri* genome and submitted to analysis for common or different motifs using the Multiple Expectation maximizations for Motif Elicitation (MEME-suite) tool (Bailey & Elkan, 1994). Nucleotide motifs were scanned for biological roles using Gene Ontology for Motifs (GOMo) tool (Buske *et al.*, 2010) to determine if any motif was significantly associated with genes linked to one or more genome ontology (Go) using *Arabidopsis thaliana* database.

**Methods S4.** Involvement of *CYP72A1182* in different HPPD-resistant *A. palmeri* populations

Initial screening was performed in the greenhouse using one and two times the label rate of mesotrione and tembotrione to confirm herbicide resistance. Ten suspected resistant populations (WR2019-274, WR2019-141, WR2019-140, WR2019-137, WR2019-199, WR2019-200, WR2019-044, WR2019-144, WR2019-198, WR2019-273), two known tembotrione-resistant (WR2013-034 and NER), and one sensitive control (IHX_3361) population were used for the following experiments.

*Whole-plant dose response*

Seeds from 13 *A. palmeri* populations were sown in plastic trays filled with soil and kept in the greenhouse at 28º C and photoperiod of 16 h light. After one week, two seedlings were transplanted into single fiber pots with dimensions of 11 × 7 × 11.5 composing one replicate. The dose response experiment consisted of 11 doses, which were 0, 1/128, 1/64, 1/32, 1/16, 1/8, 1/4, 1/2, 1, 2 and 4 × the label rate (91 g a.i. $ha^{-1}$). Every dose had six replicates, with a total of 12 plants for each dose. The herbicide tembotrione (Laudis, 419 g a.i. $L^{-1}$, Bayer, Leverkusen, Germany) was applied together with 2,200 g a.i. $ha^{-1}$ of the wetting agent Mero (Bayer, Leverkusen, Germany) and 170 g a.i. $ha^{-1}$ ammonium sulfate using a stationary research sprayer (Höchst AG, Höchst, Germany) calibrated to deliver a spray volume of 300 L $ha^{-1}$. Survival and fresh shoot weight were recorded 28 d after application.

*$^{14}$C-Tembotrione metabolism*

Metabolism of tembotrione was measured in 13 populations over time in six individuals per population and treatment. Tembotrione was applied on the two youngest expanded leaves of individuals at the four-leaf stage with a total of ten 1-µL droplets (5 µL per leaf) of $^{14}C$-tembotrione (Bayer, Leverkusen, Germany) in a 0.3% v/v Mero solution (Bayer) with 3.3 kBq or 200,000 dpm $\mu L^{-1}$, corresponding to 0.762 µg $\mu L^{-1}$of tembotrione. Treated plants were kept in a growth chamber at 28°C under continuous light conditions with a light intensity of 500 µmol $m^{-2}$ $s^{-1}$ and 70% humidity. Plant shoots were harvested at 6, 12, 24, and 48 HAT. The harvested tissue was washed in 80% acetone three times to remove any non-absorbed $^{14}C$-tembotrione, and then disrupted in 500 µL of methanol with 5-mm stainless steel beads at 30 Hz for 10 min. The homogenate was centrifuged at 6,000 × g for 10 min. The residue was re-extracted with 600 µL of methanol followed by a final extraction with 600 µL of 90% acetonitrile. All solvents used were high-performance liquid chromatography (HPLC) grade (Sigma-Aldrich, Steinheim, Germany; ≥ 99.9 % HPLC grade). The pooled supernatant was evaporated under continuous air flow at 55 °C and then re-suspended in 200 µL of 90% acetonitrile using a shaker and ultrasonic bath and then filtered through a 0.45-µm low-binding hydrophilic polytetrafluoroethylene (PTFE) mesh for 10 min at 2,200 × g in the centrifuge. The recovered radioactivity in the filtrate was 92% of the total applied, on average. A non-treated control sample, spiked with $^{14}C$-tembotrione just prior to extraction, was also included. Separation and HPLC identification of the parent tembotrione herbicide and its metabolites were performed on a reverse-phase HPLC system (LC Net II/ADC with PU-980 pump unit, LC-980-02 gradient unit and CO-2060 Plus column thermostat; Jasco, Oklahoma City, OK, USA). Chromatographic separation was achieved with a 150 × 2.0 or 3.0 mm internal diameter (I.D.) Luna C18 (2) column with a particle size of 3 µm (Phenonemex, Aschaffenburg, Germany) at a flow rate of 0.5 mL $min^{-1}$. The mobile phases consisted of 0.05% phosphoric acid (A) and acetonitrile: 0.05% formic acid (B) and were run at a

60-min linear gradient from 0 to 60% solvent B, followed by a 1-min linear gradient from 60 to 90% solvent B, plateauing for 4 min. The column was then flushed with 100% solvent A for 7 min.

*P450 gene expression*

To ascertain the involvement of *CYP72A1182* in the newly identified resistant populations, a gene expression experiment was conducted. The aim was to assess the expression levels of *CYP72A1182* and determine its potential role in conferring resistance within the populations. Plant growth and herbicide application at 91 g a.i. $ha^{-1}$ were performed as described previously. The Nebraska sensitive population (Küpper *et al.*, 2018) was used as a second negative control in this experiment. Four plants of each population were used for gene expression. Tissue of the first and second youngest leaves were harvested before and 6 HAT, respectively. Tissue was collected into 2 mL Eppendorf tubes and placed in liquid nitrogen. The tubes were kept in -80 ºC for further analysis. Tissue was ground with 3 mm stainless steel beads in a TissueLyser (Qiagen) with intensity of 30 for 1 min. RNA was isolated using Direct-zol RNA Miniprep from Zymo Research and purified with DNase I. cDNA synthesis was performed using ProtoScript® II First Strand cDNA synthesis kit using 1 µg of RNA. Gene expression analysis and conditions were the same as defined in the section of gene validation. The reference gene used in the experiment was *18S rRNA*, whereas the candidate genes tested were *CYP72A1182* and *CYP81CJ2*.

*Gene copy number*

The six *A. palmeri* populations most resistant to tembotrione described above, along with NES, NER, and another sensitive population IHX_3361, were used for *CYP72A1182* gene copy number analysis. Plant growth, tissue collection, and DNA extraction were performed as described. Genomic DNA at 50 ng/µL was used for relative gene copy quantification using a modified method $2^{-\Delta\Delta ct}$ (Livak & Schmittgen, 2001). The *ALS* gene was used as a single-copy control gene. The primers for *ALS* were used from previous research studying *EPSPS* copy number in *A. palmeri* (Gaines *et al.*, 2010). Relative quantification of *CYP72A1027* was calculated with a modified method ΔCt (Ct, ALS – Ct, CYP). qPCR conditions were the same as described in the section for gene expression validation. Each population had eight biological samples run in two technical replicates.

**Methods S5.** Dose-response and segregation analysis

A dose-response was conducted to define a delimiting rate to best differentiate the S × S (NES) and R × R (NER) populations. Seed of NES, NER, and two F1 populations NERmale × NESfemale (crosses A and B) were sown on soil and germinated in the greenhouse at 28º C and 16h light photoperiod then transplanted to 4 × 4 cm inserts after 7 d. The dose response experiment consisted of 11 doses, which were 0, 1/128, 1/64, 1/32, 1/16, 1/8, 1/4, 1/2, 1, 2 and 4 × the label rate (91 g a.i. $ha^{-1}$). Each dose was applied on two plants with eight replicates. The herbicide tembotrione (Laudis, 419 g a.i. $L^{-1}$, Bayer, Leverkusen, Germany) was applied together with 1% v/v MSO using an automated spray chamber (Greenhouse Spray Chamber, model Generation IV) using a TJ8002E nozzle, calibrated to deliver 200 L $ha^{-1}$ at a pressure of 280 kPa and speed of 1.2 m $s^{-1}$. Survival and fresh shoot mass were recorded 28 d after application and fitted to a three-parameter log-logistic model using the drc package in R (Ritz *et al.*, 2015).

Segregation in the pseudo-F2 population was performed in response to a delimiting rate of 77 g $ha^{-1}$ with 606 plants in cross A and 1,277 plants in cross B. At 28 d after tembotrione application, plants were rated and shoot fresh mass was measured. For QTL analysis, the 71 and 110 most susceptible plants and 49 and 91 most resistant plants, from cross A and B, respectively, were selected based on survival and shoot fresh mass. Parental NES and NER were grown and submitted to the same herbicide screen and 20 of each population were selected for QTL mapping.

**Methods S6.** Statistical analysis

The alignment of the RNA-sequencing experiment was used to analyze differentially expressed genes (DEGs) using DEseq2 (v1.20.0) (Love *et al.*, 2014) package in R (v4.0.4) (R Core Team, 2023). Populations S and R were considered as factor of conditions. Samples from cross A and cross B were pooled together for the analysis. Contrast analyses performed were R versus S before treatment (0 HAT), and 6 and 12 HAT, and 6 HAT versus 0 HAT and 12 HAT versus 0 HAT for each population. The data were submitted for shrinkage log2 foldchange (LFC). Counts were normalized by creating a “virtual reference sample” using geometric mean of counts over all samples for each gene (Anders & Huber, 2010). Principal component analysis (PCA), volcano and MA plots were performed on the normalized gene expression data. Genes with less than ten reads were excluded from analysis. A P-adjusted value $<0.05$ cutoff and log2 fold-change of 1 was used to identify DEGs.

The validation of candidate P450 genes in NER and NES was performed in four biological samples and two technical replications. Analysis of the *CYP72A1182* and *CYP81CJ2* role in the field collected *A. palmeri* populations from 2019 were performed using four biological samples and two technical replications. The mean Ct values and the standard deviation were calculated by treatment. A melt curve analysis confirmed the presence of a single amplified product for each reaction, based on presence of a single melting temperature consistent across samples for each gene. Relative transcript abundance was calculated using $2^{(-\Delta\Delta Ct)}$ method (Livak & Schmittgen, 2001). The reference population used for the P450 validation analysis was NES untreated. To analyze the role of P450 genes in the field collected populations, the sensitive control (IHX_3361) before herbicide application was used as control. Fisher's LSD's test ($p < 0.05$) was used to compare the relative expression between treatments for each experiment.

For the whole-plant dose responses, the statistical software R v.3.5.3 (Ritz *et al.*, 2015) was used for data analysis. Data were adjusted using the three-parameter log-logistic model with the function modelFit () from the drc package (Ritz *et al.*, 2015). The herbicide doses that caused a 50% reduction in each variable were estimated using the model: $Y=d/1+\exp[b(\log x-\log e)]$, where $d$ is the upper limit, $b$ is the slope, $x$ is the dose, and $e$ is the dose that causes 50% reduction in Y. The statistical difference between the resistant biotype and the sensitive biotype for $ED_{50}$ (survival) or $GR_{50}$ (fresh shoot weight) was calculated using the function EDcomp (). Resistance index (RI) was calculated using the ratio of $GR_{50}$ values of each biotype with the sensitive biotype. Graphs were generated using GraphPad Prism version 8.2.1 (San Diego, California).

For $^{14}C$-tembotrione metabolism analysis, the identification of the main metabolites M1, M2, M3, M4, and M5 were based on previous results (Küpper *et al.*, 2018) and retention time. Peak area (% of recovered) was obtained from HPLC and used to quantify and compare metabolites. The means of six replications of each biotype were compared to IHX_3361 (sensitive) population by Dunnett's multiple comparison.

**Notes S1** Alignment of *CYP81CJ2* and *CYP72A1182* promoter between susceptible and resistant *Amaranthus palmeri*.

### Alignment of *CYP81CJ2* promoter of S and R plant

```
#=======================================
#
# Aligned_sequences: 2
# 1: S1
# 2: R1
# Matrix: EDNAFULL
# Gap_penalty: 10.0
# Extend_penalty: 0.5
#
# Length: 1250
# Identity:    1250/1250 (100.0%)
# Similarity:  1250/1250 (100.0%)
# Gaps:           0/1250 ( 0.0%)
# Score: 6250.0
#
#
#=======================================

S1                 1 AAAATACTTCTATAAACTCTTAACACTACCATAAACCAAACCCTACTTTA     50
                     ||||||||||||||||||||||||||||||||||||||||||||||||||
R1                 1 AAAATACTTCTATAAACTCTTAACACTACCATAAACCAAACCCTACTTTA     50

S1                51 CCTACATACCACTCACATAAATATTACGATCTATTTTATTAAGTAAAGAA    100
                     ||||||||||||||||||||||||||||||||||||||||||||||||||
R1                51 CCTACATACCACTCACATAAATATTACGATCTATTTTATTAAGTAAAGAA    100

S1               101 TTCTTTGTTTGTTCCTAAAGAAAACTATAAATTAAATACCTATACTCTCT    150
                     ||||||||||||||||||||||||||||||||||||||||||||||||||
R1               101 TTCTTTGTTTGTTCCTAAAGAAAACTATAAATTAAATACCTATACTCTCT    150

S1               151 AAATAACTTTATCTACCATGTGATGGGTGTACCTACGTACATCGGAAAGA    200
                     ||||||||||||||||||||||||||||||||||||||||||||||||||
R1               151 AAATAACTTTATCTACCATGTGATGGGTGTACCTACGTACATCGGAAAGA    200

S1               201 AAATCTTATCTCATTTACCATTCATCTTTCTCAGTTTAACTCAAATTTAT    250
                     ||||||||||||||||||||||||||||||||||||||||||||||||||
R1               201 AAATCTTATCTCATTTACCATTCATCTTTCTCAGTTTAACTCAAATTTAT    250

S1               251 ACCTATTTACAGAAGTAGTTTAACGTGAATATAACTAATAAGTATCCTTA    300
                     ||||||||||||||||||||||||||||||||||||||||||||||||||
R1               251 ACCTATTTACAGAAGTAGTTTAACGTGAATATAACTAATAAGTATCCTTA    300

S1               301 TACACGTTAATTAAATCTAGACAAATACCCGCCCAATTAGTCTCAGACAG    350
                     ||||||||||||||||||||||||||||||||||||||||||||||||||
R1               301 TACACGTTAATTAAATCTAGACAAATACCCGCCCAATTAGTCTCAGACAG    350

S1               351 GTAAATTAATCCTCTTATCCACTTATTTTTATCTTAAAAATTTCATAATT    400
                     ||||||||||||||||||||||||||||||||||||||||||||||||||
R1               351 GTAAATTAATCCTCTTATCCACTTATTTTTATCTTAAAAATTTCATAATT    400

S1               401 AATCAAAAAACTAACGAATTCTAACACATGAAATCACTCCATAACATCCT    450
                     ||||||||||||||||||||||||||||||||||||||||||||||||||
R1               401 AATCAAAAAACTAACGAATTCTAACACATGAAATCACTCCATAACATCCT    450

S1               451 CCACAATACATACATGAAATTAATATCTCATTAATATCCAAAATTATATC    500
                     ||||||||||||||||||||||||||||||||||||||||||||||||||
R1               451 CCACAATACATACATGAAATTAATATCTCATTAATATCCAAAATTATATC    500
```

```
S1              501 TTTTCTACTTATTGAAAATCCAAAAACCCTCTATAATATTCATTTAAATA    550
                    ||||||||||||||||||||||||||||||||||||||||||||||||||
R1              501 TTTTCTACTTATTGAAAATCCAAAAACCCTCTATAATATTCATTTAAATA    550

S1              551 CATACTTTCCCATTACCCCGTTTAATATACCTGTTTGATTTTAACATATA    600
                    ||||||||||||||||||||||||||||||||||||||||||||||||||
R1              551 CATACTTTCCCATTACCCCGTTTAATATACCTGTTTGATTTTAACATATA    600

S1              601 TCTCGTTTTAATTTCTCTAGTGAATTAAATCCTCTATAAAAGAGTGCATT    650
                    ||||||||||||||||||||||||||||||||||||||||||||||||||
R1              601 TCTCGTTTTAATTTCTCTAGTGAATTAAATCCTCTATAAAAGAGTGCATT    650

S1              651 TTTAGTACACAATTTTTACACTTACGTTTAGTACAGTAAGAGTATGTGCT    700
                    ||||||||||||||||||||||||||||||||||||||||||||||||||
R1              651 TTTAGTACACAATTTTTACACTTACGTTTAGTACAGTAAGAGTATGTGCT    700

S1              701 AAAGTATAAAAAGTGCATTTTATTAATAATTCCAACAGAAGATTATAGTA    750
                    ||||||||||||||||||||||||||||||||||||||||||||||||||
R1              701 AAAGTATAAAAAGTGCATTTTATTAATAATTCCAACAGAAGATTATAGTA    750

S1              751 TAATTCTTTGATTAATTAATCGGTACTCAAAGTCAAATACCAACATTGAG    800
                    ||||||||||||||||||||||||||||||||||||||||||||||||||
R1              751 TAATTCTTTGATTAATTAATCGGTACTCAAAGTCAAATACCAACATTGAG    800

S1              801 ACTATACAAATATAATAGAGTTTGAATATTTGAGATCAGGTACTATACGA    850
                    ||||||||||||||||||||||||||||||||||||||||||||||||||
R1              801 ACTATACAAATATAATAGAGTTTGAATATTTGAGATCAGGTACTATACGA    850

S1              851 TTATAAGCTAGTTCAAATATATCAAGTTAATATTCCTCAAAAGTATATGA    900
                    ||||||||||||||||||||||||||||||||||||||||||||||||||
R1              851 TTATAAGCTAGTTCAAATATATCAAGTTAATATTCCTCAAAAGTATATGA    900

S1              901 CATAACGAATTAAATCGATGAGAATACTCTCTAGCAGAAAATTTCTATGT    950
                    ||||||||||||||||||||||||||||||||||||||||||||||||||
R1              901 CATAACGAATTAAATCGATGAGAATACTCTCTAGCAGAAAATTTCTATGT    950

S1              951 TAAGTATTTATTATTCAGGTGTAAGAATAAAAGAGAAATTAAACATAATT   1000
                    ||||||||||||||||||||||||||||||||||||||||||||||||||
R1              951 TAAGTATTTATTATTCAGGTGTAAGAATAAAAGAGAAATTAAACATAATT   1000

S1             1001 CACCCGGTAAATCGGGTATTTATATTATAACACAGAGAGCCTCTCTGGCA   1050
                    ||||||||||||||||||||||||||||||||||||||||||||||||||
R1             1001 CACCCGGTAAATCGGGTATTTATATTATAACACAGAGAGCCTCTCTGGCA   1050

S1             1051 GCGAGTGTTGTTAAACACCATAACGATATAATAGTTGTTTCTGTAATTAT   1100
                    ||||||||||||||||||||||||||||||||||||||||||||||||||
R1             1051 GCGAGTGTTGTTAAACACCATAACGATATAATAGTTGTTTCTGTAATTAT   1100

S1             1101 AACTATATCATTATGTTGTCTTAATATTTTTCTCTTGTTGAATTATAAAA   1150
                    ||||||||||||||||||||||||||||||||||||||||||||||||||
R1             1101 AACTATATCATTATGTTGTCTTAATATTTTTCTCTTGTTGAATTATAAAA   1150

S1             1151 TACTTTTTTATACTCTTTATATTTTTTCATTTTTTTTAGTTTATATCAAT   1200
                    ||||||||||||||||||||||||||||||||||||||||||||||||||
R1             1151 TACTTTTTTATACTCTTTATATTTTTTCATTTTTTTTAGTTTATATCAAT   1200

S1             1201 TTTATATTCATCGTTTAAATTATCTTGTCACTTTTTTTAACTTTATTTCA   1250
                    ||||||||||||||||||||||||||||||||||||||||||||||||||
R1             1201 TTTATATTCATCGTTTAAATTATCTTGTCACTTTTTTTAACTTTATTTCA   1250

#---------------------------------------
#---------------------------------------
```

**Alignment of *CYP72A1182* promoter of S and R plant**

```
#=======================================
#
# Aligned_sequences: 2
# 1: S1
# 2: R1
# Matrix: EDNAFULL
# Gap_penalty: 10.0
# Extend_penalty: 0.5
#
# Length: 2072
# Identity:    1593/2072 (76.9%)
# Similarity:  1593/2072 (76.9%)
# Gaps:         382/2072 (18.4%)
# Score: 7172.5
#
#
#=======================================

S1                 1 AATATATAAAACAAAATAACTCTTTACCTAATAATTAAGGATTA-ATATA     49
                     |||||||||||.|||||||||||||||||||||||||||||||| |||||
R1                 1 AATATATAAAATAAAATAACTCTTTACCTAATAATTAAGGATTACATATA     50

S1                50 TATCTCGAAAACAACATTTACTTCCCTATCCCTATATGTGATACACACCT     99
                     |||||||||||||||||||||||||||||||||             ||||
R1                51 TATCTCGAAAACAACATTTACTTCCCTATCCCT-------------ACCT     87

S1               100 TCACTTTGTTTTCTTCCTTTACTCTCATATATATCCTATTGATTTTTTGT    149
                     ||||||||||||||||||||||||||||||||||||.|||||||||||||
R1                88 TCACTTTGTTTTCTTCCTTTACTCTCATATATATCCGATTGATTTTTTGT    137

S1               150 AGTAGTAGTTGGGTTATAAGGCGAATTTTTGCCCCTACCTTTTCTTGTTG    199
                     ||||||||||.||||||||||||||||||||||.||||||||||||||||
R1               138 AGTAGTAGTTAGGTTATAAGGCGAATTTTTGCCTCTACCTTTTCTTGTTG    187

S1               200 TATATCTTAAAATACTCACTTATTCTCAGTTCTTTTACTACAACTATTAA    249
                     ||||||||||||||||||||||||||||||||||||||||||||||||||
R1               188 TATATCTTAAAATACTCACTTATTCTCAGTTCTTTTACTACAACTATTAA    237

S1               250 CTTCTTATAATACACCTATCTCTTGTTTGGAATGTTATTTATCTTGAGTA    299
                     ||||||||||||||||||||||||||||||||||||||||||||||||||
R1               238 CTTCTTATAATACACCTATCTCTTGTTTGGAATGTTATTTATCTTGAGTA    287

S1               300 GTAGTAGTATAGGTCACATAGGGCGAGTATCTATTTGATACTAGTCTCAA    349
                     ||||||||||          ||||.|||||||||||||||||||||||||
R1               288 GTAGTAGTAT----------GGGCTAGTATCTATTTGATACTAGTCTCAA    327

S1               350 ACCCCTCCCTTCTATC-----TTGAGTATCGGTACATCCTCTCATGCTAG    394
                     ||||||||||| ||||     |.||||||.||||||||||||.|||||||
R1               328 ACCCCTCCCTT-TATCTGCCGTCGAGTATGGGTACATCCTCTTATGCTAG    376

S1               395 TCTCTCAGGGGGCCGAGGCATACAAGAAGTTTCACTCTCC---ATGGACG    441
                     |||||.| ||||||||||.||||||||||||||.||||||   ||.|.|.
R1               377 TCTCTTA-GGGGCCGAGGTATACAAGAAGTTTCTCTCTCCATAATCGTCA    425

S1               442 TTAATAAACTTTCGAACGTTCTGTGTGGGT--------GCAATTGTTACG    483
                     ||||||.|||||||...|||||||||||||        ||||||||||||
R1               426 TTAATAGACTTTCGGTTGTTCTGTGTGGGTGCAATTCAGCAATTGTTACG    475

S1               484 TATCATTAGCTAATTGATTTCTATGTGTTGTTTTTCTTATAATGATTATT    533
                     |||||||||||||||||||||||||||||||||||||||| |||||||||
R1               476 TATCATTAGCTAATTGATTTCTATGTGTTGTTTTTCTTAT-ATGATTATT    524
```

```
S1          534 TATCTTGATATTTTCATATAAAACTTGCCTAAGCCGCGTTCCATCAACAT   583
                ||||||||||||||||||||||||||||||||||||||||||||||||||
R1          525 TATCTTGATATTTTCATATAAAACTTGCCTAAGCCGCGTTCCATCAACAT   574

S1          584 ACTCATCAACAATTGACAGATTTATCGATAATCAACAGCTATCAACAACA   633
                ||||||||||||||||||||||||||||||||||||||||||||||||||
R1          575 ACTCATCAACAATTGACAGATTTATCGATAATCAACAGCTATCAACAACA   624

S1          634 TTTTAATACCAATAATCTATACATCTAGTGAAAAATTAACACATTTTATA   683
                ||||||||||||||||||||||||||||||||||||||||||||||||||
R1          625 TTTTAATACCAATAATCTATACATCTAGTGAAAAATTAACACATTTTATA   674

S1          684 ATAATAAACCATTTTCACCGCGAGTTCCATTTACAATGAACAATCTTACA   733
                |||||||||||||||||.|||||.||||||||||.|||||||||.|||||
R1          675 ATAATAAACCATTTTCATCGCGAATTCCATTTACGATGAACAATTTTACA   724

S1          734 TTAGAATAACTCTTTTTGTGTTTTAGAGTGTTCTCTGCCAGATATCCTAT   783
                || |||||||||||
R1          725 TT-GAATAACTCTT------------------------------------   737

S1          784 CTGGTGTTTTAACCCGGTCGGGTAAAGGGAATTGACTAAAAAAATAATAT   833
                                                          ||||
R1          738 ------------------------------------------AATA----   741

S1          834 TATAGTTAATATAATACAATATTAGAATTTACATTCCATAATATTGGAGC   883

R1          742 --------------------------------------------------   741

S1          884 TTATATGATAAGATATTGGGGTTTATATTCAATAATATCGGGGTTTATTA   933

R1          742 --------------------------------------------------   741

S1          934 AATAATATTGGGGTTTATATGATACAATATAAAACCCGACCCGAAATACC   983

R1          742 --------------------------------------------------   741

S1          984 TAAGGCAGATATCTTATCTGGCAGAGGGTACTCTGAATGACACTCTTTTT  1033
                                             ||||             ||||
R1          742 -----------------------------ACTC-------------TTTT   749

S1         1034 TATATCTCTTTTAAGAGTTTTAGAATAAACCTATGATGAA-TTTTACAAT  1082
                ||||||||||||||||||||||.||||||..|.||||||| |||||||||
R1          750 TATATCTCTTTTAAGAGTTTTACAATAAATTTTTGATGAATTTTTACAAT   799

S1         1083 GAAATAGCTTTGTACTATAAAAAGAATTCACAAAATCGGAGTAATTTAGT  1132
                ||||||||.||||||.||||||||||||||||||||||||||||||||||
R1          800 GAAATAGCATTGTACCATAAAAAGAATTCACAAAATCGGAGTAATTTAGT   849

S1         1133 AATTGACAGCAAACCGTAAGCTGTAAATTACTATACTTTCTATAACTTTT  1182
                |||.|||||||||||.|||.||.|||||||||||||||.|||||||||||
R1          850 AATCGACAGCAAACCATAACCTATAAATTACTATACTTACTATAACTTTT   899

S1         1183 AAAGA--ATATTAAACCCACCTTTTTTGAGAACTGTTCAAATTACGAGAA  1230
                ||| |  |||||||||||||||||||||||||||.|.|||||||||||||
R1          900 AAA-ATCATATTAAACCCACCTTTTTTGAGAACTTTACAAATTACGAGAA   948

S1         1231 TAAATAAAATGAAGTTAGCAGAATAAAAGAGATGTTTTAAGTAAAAGTGC  1280
                ||||.|||||||||||||||||.|||||||||||||||||.|||.|||||
R1          949 TAAACAAAATGAAGTTAGCAGAGTAAAAGAGATGTTTTAATTAAGAGTGC   998

S1         1281 GTAATAGTAAATTTTTACTCTGTAATCTATCAACTATTCGTAATGAAATA  1330
                |||||||||||.|||||.|||.||||||||||..||||||||||||||||
```

```
R1            999 GTAATAGTAAACTTTTATTCTATAATCTATCACGTATTCGTAATGAAATA   1048

S1           1331 TACGTTGTATTCCCATCT-AAAAAAGAAATGATGTGGTTTATAAGTAATA   1379
                  |||||||||||.|||||| |||||||||||.||||||||||||||||||
R1           1049 TACGTTGTATTGCCATCTAAAAAAAGAAATAATGTGGTTTATAAGTAATA   1098

S1           1380 CTAAAAATCGGGAAAATAATTTATAAGAAATCTTCCTAAAGAAGCTAATT   1429
                  ||||||||||||||||||||||||||.|.|||||||||||||.||||||
R1           1099 CTAAAAATCGGGAAAATAATTTATAAAACATCTTCCTAAAGAAACTAATT   1148

S1           1430 TTACATTGTAGATCTACTTAATACTAGAGAATATTTAATTAATTATATAT   1479
                  |||.||||||.|||.||||||||||||||||||||||||||||||||.|
R1           1149 TTAAATTGTAAATCAACTTAATACTAGAGAATATTTAATTAATTATATCT   1198

S1           1480 AAGTTACATTTAATCCGAAATTTAGTACTACTAAATATGATAAT------   1523
                  ||||||||||||||||||||||||||||.||||||||||||||
R1           1199 AAGTTACATTTAATCCGAAATTTAGTACTGCTAAATATGATAATGTACTA   1248

S1           1524 -------AATATTAACAGAATTAAAATGCGATTCAATGATTTAT--TATA   1564
                         |||||||||.||||||||.|||||||||||||||||  ||||
R1           1249 ATATTTAAATATTAACGGAATTAAAGTGCGATTCAATGATTTATTGTATA   1298

S1           1565 A-------------------------------ATTACTCTATAATGTTAC   1583
                  |                               ||||||||||||||||||
R1           1299 AAATAGAGTTTTTTTTCAATGATTTATTGTATATTACTCTATAATGTTAC   1348

S1           1584 TAGCTTATTATTTAACATCTTCTGTACATTTTACTATGATTATGGTTATA   1633
                  ||||||||||||||||||||||||||.||||||||||..|||||||||||
R1           1349 TAGCTTATTATTTAACATCTTCTGTAAATTTTACTATACTTATGGTTATA   1398

S1           1634 TGAAGGAGGCAAAATGATATGAATGATGTAAAATGAAAATGCGTTAAGTT   1683
                  |||||||.||||..|.|||||||.||||||||||.|||||||||||.|||
R1           1399 TGAAGGAAGCAAGGTCATATGAACGATGTAAAATAAAAATGCGTTAGGTT   1448

S1           1684 ACGCAATAAAGTTAGTAAATATATAACTTAATATGTGTAGATTTTTAATA   1733
                  |||.||||||||||||||||||.|||.|||||||||..||||||.|||||
R1           1449 ACGTAATAAAGTTAGTAAATATCTAATTTAATATGTACAGATTTATAATA   1498

S1           1734 TTTTTCATTTATACTAATTTCATCCGCTAATCTGAAAAATTTGTTCTTAG   1783
                  |||||||.||||.|||..|||||||.||||||||..||.||||||||||.
R1           1499 TTTTTCAATTATCCTATATTCATCCACTAATCTGCTAAGTTTGTTCTTAA   1548

S1           1784 GTGAACTGATATAAAAAAGAATGTGTAATCGGTGTATATTTTATTCTAAT   1833
                  |||||||||||||||||||||||||||||..||||||||||||||..|||
R1           1549 GTGAACTGATATAAAAAAGAATGTGTAATTAGTGTATATTTTATTTGAAT   1598

S1           1834 TTGCTACTTATCATACTTATTATCTTTACATCGTTCATAAAGTCTAACCT   1883
                  ||..||||||||||.|.|||||||.|||||||||.||||||||||| .|||
R1           1599 TTATTACTTATCACAGTTATTATTTTTACATCGCTCATAAAGTCT-GCCT   1647

S1           1884 TCTTAAGAGTCAGAGTAAATAAAACATACTACCTCTTTTTTTCCTACTTT   1933
                  .|||.| .|.||||||||.|||||||||||||||||||||||||||||||
R1           1648 CCTTCA-TGGCAGAGTAATTAAAACATACTACCTCTTTTTTTCCTACTTT   1696

S1           1934 TAACTTATCATACACCACTAATCATTATTCACCTCTACTAAACCC-AATT   1982
                  ||||||||||||||||||||||||||||||||||||||||||||| ||||
R1           1697 TAACTTATCATACACCACTAATCATTATTCACCTCTACTAAACCCAAATT   1746

S1           1983 CTGTTCTTTACT----------   1994
                  ||||||||||||
R1           1747 CTGTTCTTTACTATTCTCTGGG   1768
```

#-------------------------------------
#-------------------------------------
